# Viral infection disrupts intestinal homeostasis via Sting-dependent NF-kB signaling

**DOI:** 10.1101/2023.11.16.567400

**Authors:** Jared C. Nigg, Hervé Blanc, Lionel Frangeul, Vanesa Mongelli, Xavier Godron, Allison J. Bardin, Maria-Carla Saleh

## Abstract

Host-microbe interactions influence intestinal stem cell (ISC) activity to modulate epithelial turnover and composition. Here we investigated the functional impacts of viral infection on intestinal homeostasis and the mechanisms by which viral infection alters ISC activity. We report that Drosophila A virus (DAV) infection disrupts intestinal homeostasis in *Drosophila* by inducing sustained ISC proliferation, resulting in intestinal dysplasia, loss of gut barrier function, and reduced lifespan. We found that additional viruses common in laboratory-reared *Drosophila* also promote ISC proliferation. The mechanism of DAV-induced ISC proliferation involves progenitor-autonomous EGFR signaling, JNK activity in enterocytes, and requires Sting-dependent NF-kB (Relish) activity. We further demonstrate that activating Sting-Relish signaling is sufficient to induce ISC proliferation, promote intestinal dysplasia, and reduce lifespan in the absence of infection. Our results reveal that viral infection can significantly disrupt intestinal physiology, highlight a novel role for Sting-Relish signaling, and support a role for viral infection in aging.

**Graphical Abstract:** 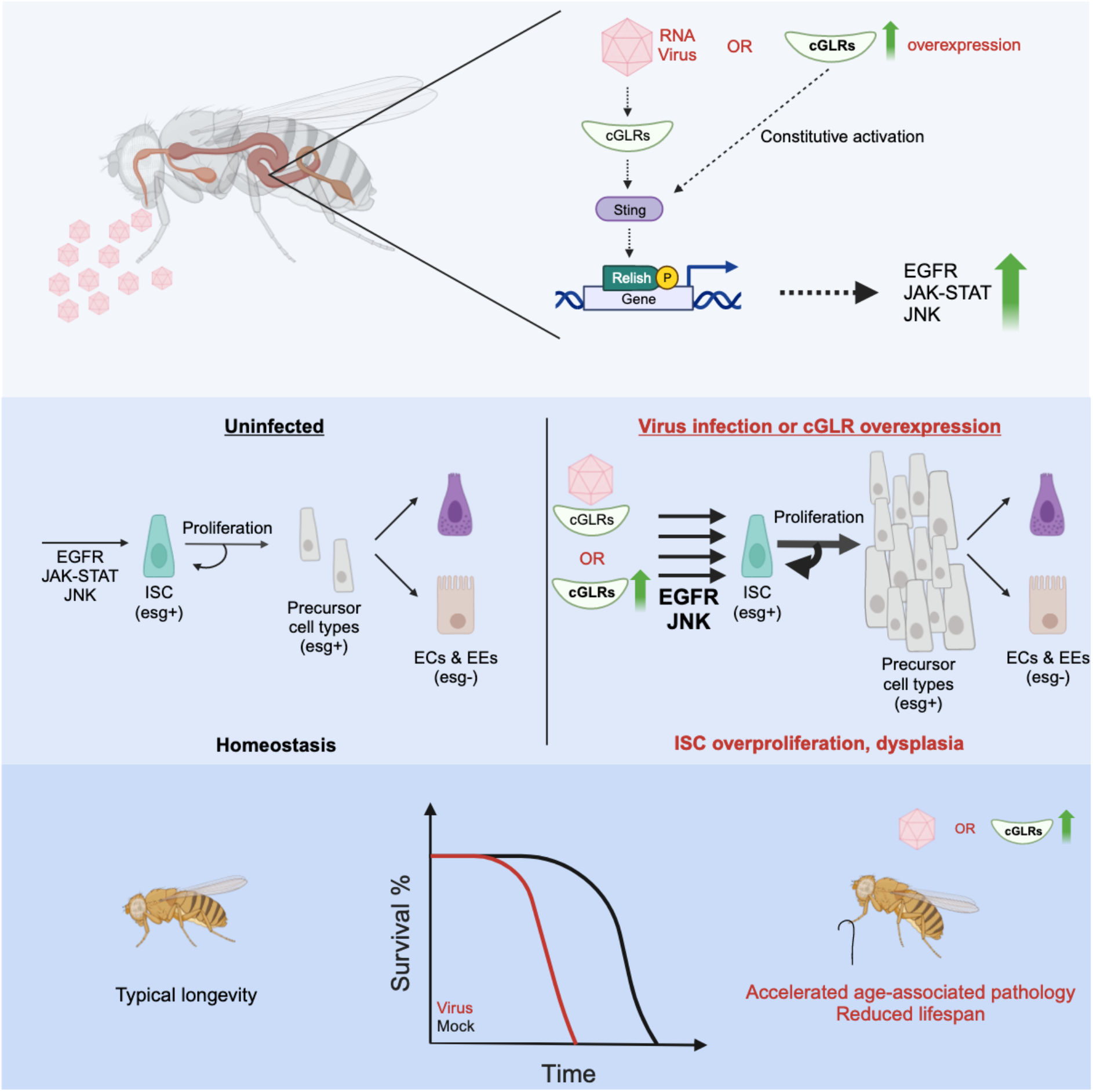

## INTRODUCTION

The gastrointestinal tract is the primary interface between internal tissues and the external environment. To maintain homeostasis in the face of constant microbial and environmental assaults, the intestinal epithelium undergoes continuous cellular turnover driven by the proliferation of intestinal stem cells (ISCs) to replace old or damaged differentiated cells^1,2^. Immune and stress signaling pathways integrate environmental cues including injury, infection, and nutrient status to modulate ISC proliferation rates, differentiated cell death, and the balance of differentiation programs in proliferating ISCs^3^. While these processes are required to replace damaged epithelial cells and maintain homeostasis, they must be tightly regulated to ensure proper intestinal function and integrity^4^. Indeed, intestinal homeostatic defects are underlying drivers of numerous conditions in mammals including inflammatory bowel disease^5^, colorectal cancer^6^, and loss of gut barrier function^7^. Consequently, maintenance of intestinal homeostasis is a primary determinant of both lifespan and healthspan, as homeostatic defects in the intestine drive inflammatory diseases and accelerate aging in the intestine and distal tissues^8–10^. Understanding how host-microbe interactions modulate intestinal homeostatic processes is thus crucial for the development of interventions to mitigate the deleterious impacts of infection and promote healthy aging.

*Drosophila melanogaster* is an experimentally tractable model at the forefront of efforts investigating how host-microbe interactions impact intestinal homeostasis. The regulatory mechanisms directing ISC proliferation in the mammalian intestine and the *D. melanogaster* midgut are highly similar^1^, both relying on several evolutionarily conserved signaling pathways, including the c-Jun N-terminal Kinase (JNK), JAK-STAT, epidermal growth factor receptor (EGFR), Wingless, and Hippo pathways. Importantly, the intestinal epithelia of mammals and *D. melanogaster* are both comprised of functionally analogous differentiated cells that are maintained by ISCs^11^. The *D. melanogaster* intestinal epithelium primarily consists of fully differentiated absorptive enterocytes (ECs) and secretory enteroendocrine cells (EEs)^12,13^. The populations of these two cell types are sustained by ISCs that proliferate and undergo partial differentiation into progenitor cells termed enteroblasts and pre-enteroendocrine cells, which themselves fully differentiate into mature ECs or EEs, respectively^12–14^.

Studies in *D. melanogaster* have revealed that pathogenic bacterial infection activates JNK signaling in damaged ECs, resulting in the release of inflammatory Unpaired (Upd) cytokines that directly and indirectly stimulate cell-autonomous JAK-STAT and EGFR signaling, respectively, in ISCs^15,16^. This triggers a transient burst of ISC proliferation that is required to repair epithelial damage following pathogen clearance^15^. In contrast, commensal bacteria stimulate low levels of ISC proliferation, promoting basal levels of epithelial renewal^15^. Age-associated shifts in the composition, abundance, and distribution of the commensal microbiota provoke chronic stress signaling that stimulates sustained over-proliferation of ISCs and disrupts homeostatic differentiation programs^15,17–21^. This process promotes intestinal dysplasia and is proposed as a primary driver of progressive age-associated gut dysfunction and mortality^17,22^. Indeed loss of gut barrier function is observed prior to and is predictive of death^4,19^. However, the precise causal relationships between dysregulated ISC proliferation, loss of gut barrier function, and death are yet uncertain.

Both chronic and acute enteric viral infections can promote intestinal pathology by disrupting intestinal homeostasis in mammals, but the molecular mechanisms underlying virus-driven homeostatic defects have not been explored in detail^23–26^. In this respect, *D. melanogaster* represents an excellent model to mechanistically investigate how host-virus interactions modulate epithelial turnover in the intestine. Natural viral infections are highly prevalent in both wild and laboratory-reared *D. melanogaster* and typically manifest as persistent infections characterized by lifelong viral replication with relatively uncharacterized impacts on host fitness and physiology^27–29^. Some of the most prevalent viruses of *D. melanogaster*, including Drosophila C virus (DCV), Nora virus, and Galbut virus^30–32^, are known to infect the intestinal epithelium, but the potential impacts of viral infection on intestinal homeostasis have not been examined.

Drosophila A virus (DAV) is a highly prevalent RNA virus of *D. melanogaster* and we previously observed that orally acquired DAV persistently infects adult flies with very high efficiency^33^. We thus focused on oral DAV infection to elucidate the potential impact of host-virus interactions on intestinal homeostasis. Here we demonstrate that DAV infection induces sustained ISC proliferation and accelerates development of age-associated intestinal pathology, thereby reducing the lifespan of infected flies. Similar observations were made during infections with other RNA viruses, suggesting that modulation of intestinal physiology is a common feature of viral infection. We found that classical epithelial repair pathways as well as novel mechanisms play roles in virus-induced ISC proliferation and the disruption of intestinal homeostasis by viral infection. Overall, our results establish the utility of *D. melanogaster* as a model for studies of host-virus interactions in the intestine, identify novel regulators of ISC proliferation, and uncover significant physiological impacts caused by highly prevalent viruses. We found that these impacts were mediated by signaling pathways that are highly conserved in humans, thus providing an excellent basis with which to further investigate intestinal host-virus interactions in health and disease.

## RESULTS

### DAV persistently infects the adult midgut

To determine if DAV infects the midgut we performed negative strand-specific RT-qPCR using RNA from midguts and carcasses dissected from orally infected flies. This assay quantifies levels of actively replicating viral RNA by detecting the negative strand of the DAV genome, which is only produced during viral replication. We detected DAV replication in midguts and carcasses by 3 days post-infection (dpi) (Figure 1A). Orally acquired DAV persistently infected both the midgut and carcass, with high levels of replicative DAV RNA observed in both tissues until at least 18 dpi (Figure 1A). Oral DAV infection was not acutely lethal. Instead, flies orally infected with DAV exhibited reduced lifespans compared to mock-infected controls at both 25 °C (Fig S1A) and 29 °C (Figure 1B). We chose to perform all subsequent experiments at 29 °C to facilitate comparison of infection phenotypes across all potential fly genotypes, including those encoding constructs for temperature-sensitive gene expression. We further note that these and all subsequent experiments were performed using female flies.

**Figure 1.**
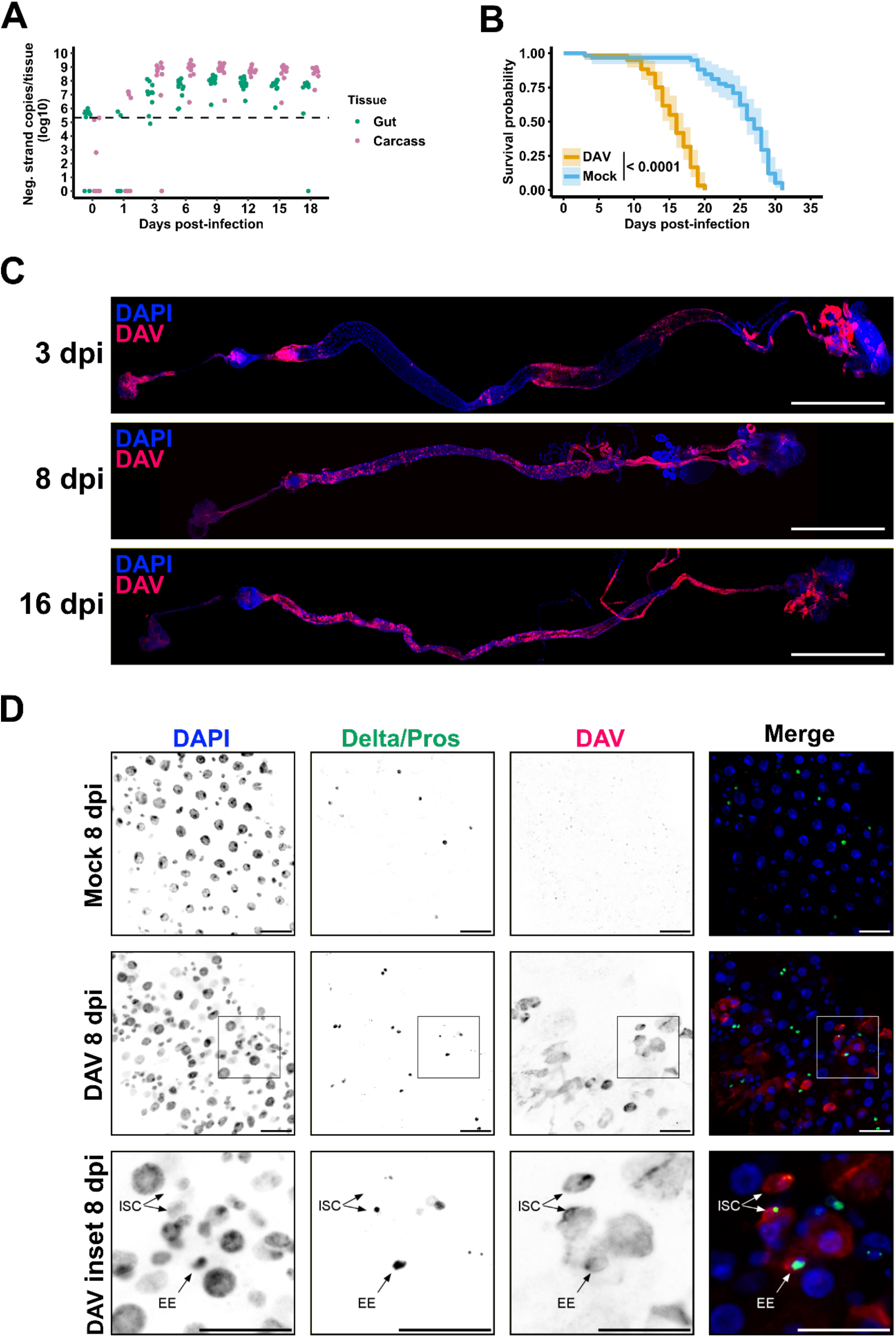
Orally-acquired DAV persistently infects the adult midgut. (A) Negative strand-specific RT-qPCR of DAV RNA in carcasses and dissected midguts from flies orally infected with DAV. (B) Survival of mock-infected and DAV-infected flies maintained at 29 °C. Shaded regions: 95% confidence intervals. Three biological replicates (n=20 flies/replicate) were analyzed. The p-value from a log-rank (Mantel-Cox) test is shown. (C) RNA FISH of positive strand DAV RNA in guts from DAV-infected flies. Scale bars: 1 mm. (D) Representative images of R4 midgut regions from mock-infected and DAV-infected flies. Boxes in the middle panels indicate the region in the lower panels. Delta+ and Pros+ cells are distinguished by membrane/vesical and nuclear staining, respectively. Arrows indicate DAV infected ISCs or EEs. Scale bars: 50 μm. See also Figures S1 and S2.

To further characterize DAV infection of the gastrointestinal tract we next determined the regional and temporal tropism of DAV infection in this tissue by detecting DAV genomic RNA by RNA FISH. At 3 dpi, a time corresponding to the onset of DAV replication, we detected DAV RNA in isolated patches throughout the alimentary canal, including the crop, foregut, cardia, all midgut regions, hindgut, rectum, and the malpighian tubules (Figures 1C and S2A). DAV RNA was evenly spread throughout the gastrointestinal tract by 8 dpi, a pattern that continued until at least 16 dpi (Figure 1C and S2A). We next determined the cellular tropism of DAV infection in the midgut by immunofluorescence with antibodies against the DAV capsid protein, Delta (an ISC marker)^34^, and Prospero (Pros; an EE marker)^34^. At 8 dpi we primarily observed infection in large Delta/Pros negative ECs, however we occasionally observed infection of Delta positive ISCs and, less frequently, Pros positive EEs (Figure 1D).

### DAV infection reduces lifespan by driving over-proliferation of ISCs

We noticed that the epithelia of midguts of DAV-infected flies exhibited clear irregularities, characterized by aberrant spatial organization and altered size distribution of nuclei compared to mock-infected midguts (Figure 1D). This phenotype resembles age-dependent intestinal dysplasia, which is driven by over-proliferation of ISCs and disruption of cellular differentiation programs due to chronic stress signaling trigged by commensal dysbiosis in aged flies^15,17,18,35^. We hypothesized that persistent DAV infection may similarly simulate chronic mitogenic signaling, leading to the premature development of intestinal dysplasia due to sustained over-proliferation of ISCs. We thus measured ISC proliferation rates in DAV- and mock-infected flies by immunofluorescence with an antibody against phosphorylated histone H3 (PH3), which is a highly-specific marker for condensed chromatin found in mitotic cells and thus selectively marks proliferating ISCs in the *D. melanogaster* midgut^13^. Indeed, DAV infection induced ISC over-proliferation in adult flies by 4 dpi and DAV-induced ISC proliferation continued for the lifetime of infected flies (Figure 2A). Importantly, germ-free (GF) flies infected with DAV exhibited premature intestinal dysplasia (Figures S1B and 2B). As the levels of DAV-induced ISC proliferation were not significantly different between conventionally reared (CR) and GF flies at 8 dpi (Figure 2B), we believe that the commensal bacteria population does not have a major impact on DAV infection. Moreover, the median survival of DAV-infected flies relative to mock-infected controls was not significantly different between CR or GF conditions (Figure S1C). We note that while infected flies were maintained at 29 °C for our experiments, DAV infection also induced ISC proliferation at 25°C (Figures S1D). Together these results indicate that DAV infection induces sustained over-proliferation of ISCs, promotes premature development of intestinal dysplasia, and reduces lifespan independent of the microbiota.

**Figure 2.**
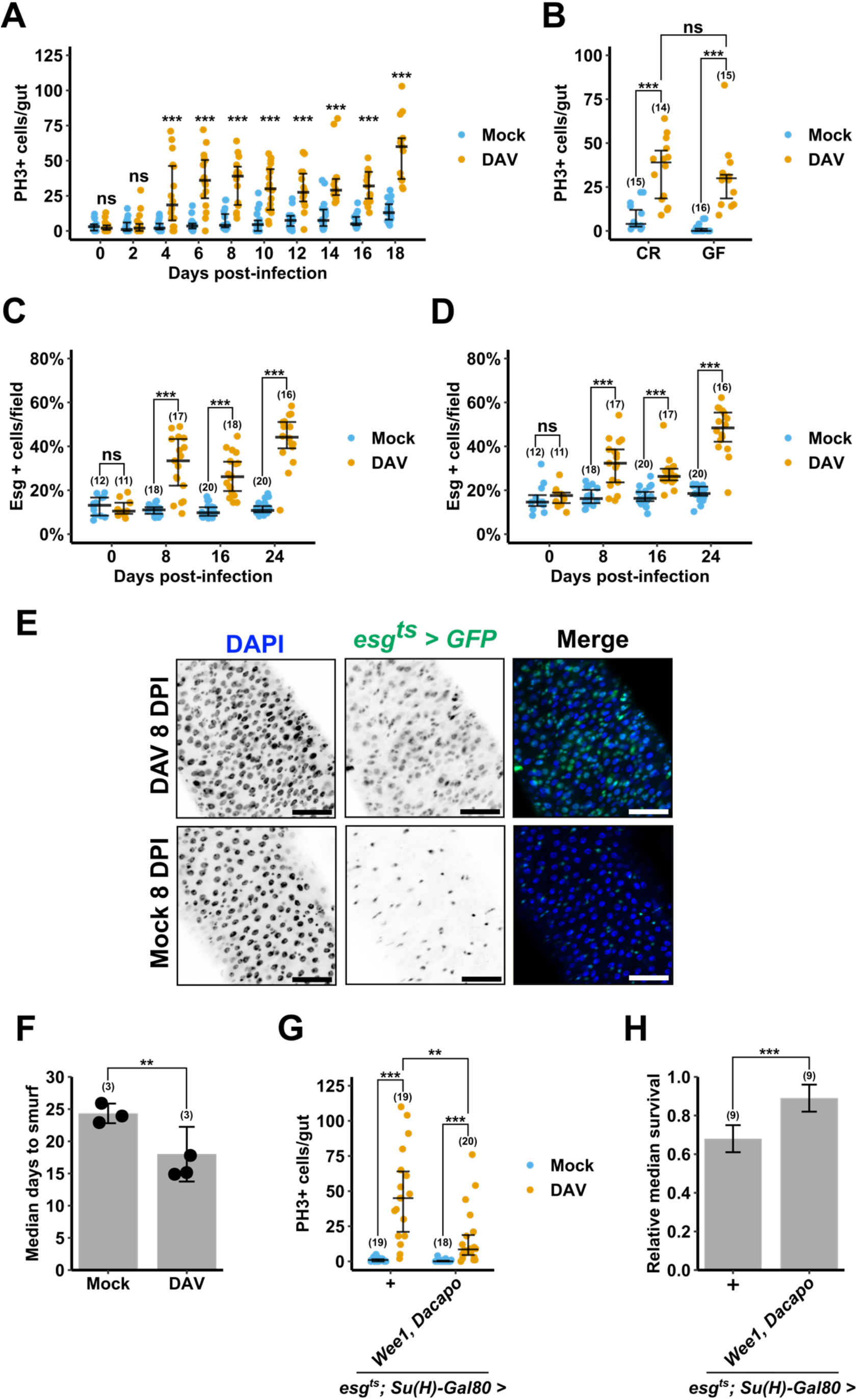
DAV infection reduces lifespan by driving over-proliferation of ISCs. (A) Quantification of PH3+ cells in midguts from mock-infected or DAV-infected flies. (B) Quantification of PH3+ cells at 8 dpi in midguts from mock-infected or DAV-infected flies maintained under CR or GF conditions. (C and D) Quantification Esg+ cells in R2 (C) and R4 (D) midgut regions from mock-infected or DAV-infected flies. (E) Representative images of Esg+ cells in R4 midgut regions from mock-infected or DAV- infected flies at 16 dpi. Scale bars: 50 μm. (F) Median days until observation of the “smurf” phenotype in mock-infected or DAV- infected flies. Dots indicate medians of three biological replicates (n=20 flies/replicate). Bar height indicates average of replicate medians. (G) Quantification of PH3+ cells at 8 dpi in midguts from mock-infected or DAV-infected flies of the indicated genotypes. (H) Relative median survival of DAV-infected flies of the indicated genotypes. Bar height indicates the average of nine biological replicates (n=20 flies/replicate) from three independent experiments. Error bars in (A-D and G) indicate median with 1^st^ and 3^rd^ quartiles. Error bars in (F and H) indicate S.D. Results were compared with a two-tailed Mann-Whitney test (A, B, and G) or a two-tailed T-test (C, D, F, and H); ns = non-significant, *p < 0.05, **p < 0.01, ***p < 0.001. Comparisons in (A) are between mock and DAV for each dpi. Numbers of biological replicates indicated in parentheses. See also Figure S1.

To determine if elevated ISC proliferation is specific to DAV, we measured ISC proliferation at 20 days post-eclosion (dpe) in flies collected from laboratory-reared stocks of isogenic wild-type (*w^1118^*) flies harboring persistent infections with either DAV, DCV, Nora virus, or Bloomfield virus (all of them RNA viruses known to infect the midgut). Persistent infections with any of the four viruses were associated with elevated ISC proliferation and irregular epithelial morphology at 20 dpe compared to uninfected controls (Figures S1E and S1F).

A hallmark of age-dependent intestinal dysplasia and loss of intestinal homeostasis is the accumulation of polyploid cells marked by their continuous expression of the progenitor cell marker, Escargot (Esg)^17,36^. We infected flies expressing GFP under the control of a temperature sensitive *esg-Gal4* driver (esg*-Gal4; tub-Gal80 UAS-GFP*^13^, referred to as *esg^ts^*) to follow accumulation of Esg+ cells during DAV infection. Consistent with our observations of altered cellular organization in DAV-infected midguts, the proportion of Esg+ cells was significantly higher in DAV-infected midguts compared to mock-infected controls in both the R2 and R4 midgut regions at 8, 16, and 24 dpi (Figures 2C-2E).

Loss of gut barrier function accompanying intestinal dysplasia is proposed as the primary driver of mortality in aged flies^4,19,37^. Given that DAV infection accelerates development of intestinal dysplasia, we used the “smurf assay^4^” to measure intestinal barrier function *in vivo* during DAV infection. In agreement with previous observations^4^, mock-infected flies showed age-dependent loss of intestinal barrier function, with 50% of mock-infected flies exhibiting the smurf phenotype by an average of 24.3 dpi (Figure 2F). DAV infection significantly reduced the age of onset of intestinal barrier dysfunction, with 50% of infected flies exhibiting the smurf phenotype by an average of 16 dpi (Figure 2F), suggesting that accelerated loss of gut barrier function may underlie the reduced lifespan of DAV-infected flies.

To more directly test a role for DAV-induced ISC proliferation in accelerating loss of intestinal homeostasis and reducing lifespan, we inhibited ISC proliferation by over-expressing the cyclin-dependent kinase inhibitors Wee1 and Dacapo in adult ISCs^38^. This was accomplished using the ISC-specific, temperature sensitive *esg-Gal4 Su(H)GBE-Gal80 tub-Gal80^ts^* system (referred to as *esg^ts^; Su(H)-Gal80*)^39^. Over-expression of Wee1 and Dacapo in ISCs did not completely block DAV-induced ISC proliferation, but did significantly reduce ISC proliferation levels compared to control flies (Figure 2G). Strikingly, reducing ISC proliferation significantly prolonged lifespan during DAV infection, as the relative median survival of DAV-infected flies was significantly greater in flies over-expressing Wee1 and Dacapo in ISCs compared to controls (Figures 2H and S1G). This result indicates that DAV- induced ISC proliferation contributes to the reduced lifespan caused by DAV infection. The prolonged lifespan accompanying inhibited ISC proliferation was not due to reduced levels of DAV replication, as DAV RNA levels were not significantly different in dissected guts or carcasses from control flies compared to those from flies over-expressing Wee1 and Dacapo in ISCs (Figures S1H and S1I). Together, our findings suggest that DAV infection reduces lifespan by accelerating the onset of classical age-associated intestinal pathologies.

### DAV infection upregulates immune, stress, and epithelial repair pathways in the intestine

We next sequenced transcriptomes from mock- and DAV-infected midguts dissected at 6 or 12 dpi to identify signaling pathways that are differentially expressed during DAV infection and which may play roles in DAV-induced ISC proliferation. We dissected midguts from both CR and GF flies to distinguish potential differential expression patterns driven independently by DAV from those that require input from the microbiota. We note that negative strand DAV RNA levels were significantly higher in midguts from GF flies than from CR flies at 12 dpi, however this difference was not observed at 6 dpi (Figure S3A).

We found that DAV infection induced significant upregulation of genes belonging to the two primary mitogenic pathways responsible for regulating ISC proliferation, the EGFR and JAK-STAT pathways (Figures 3A and 3B). Cell-autonomous EGFR signaling is required in ISCs for their proliferation and this pathway is significantly upregulated in all contexts associated with elevated ISC mitosis^40–43^. Accordingly, we found that several genes encoding components of the EGFR pathway were significantly upregulated in DAV-infected midguts at both 6 and 12 dpi in CR and GF flies (Figure 3A). While not strictly required for ISC proliferation, the JAK-STAT pathway nevertheless plays an important role in regulating ISC proliferation under both steady-state and stress-induced conditions^15,16,44^. Most notably, stressed ECs secrete Upd2 and Upd3, which activate JAK-STAT signaling in neighboring ISCs and visceral muscle cells^16,45,46^. ISC-autonomous JAK-STAT signaling is sufficient to induce ISC proliferation^16,44^. In parallel, JAK-STAT signaling in visceral muscle cells triggers secretion of the EGFR ligand Vein (Vn), which subsequently activates proliferative EGFR signaling in ISCs^40–42,45^. We found that DAV infection significantly upregulated the expression of *Upd2* and *Upd3* as well as several other JAK-STAT pathway genes in the midgut (Figure 3B). Comparison of gene expression levels in midguts from DAV-infected CR flies to midguts from DAV-infected GF flies revealed that only one EGFR- or JAK-STAT-related gene was differentially expressed between these two conditions (Figures 3A & 3B), suggesting that DAV infection activates EGFR and JAK-STAT signaling in a microbiota-independent manner.

**Figure 3.**
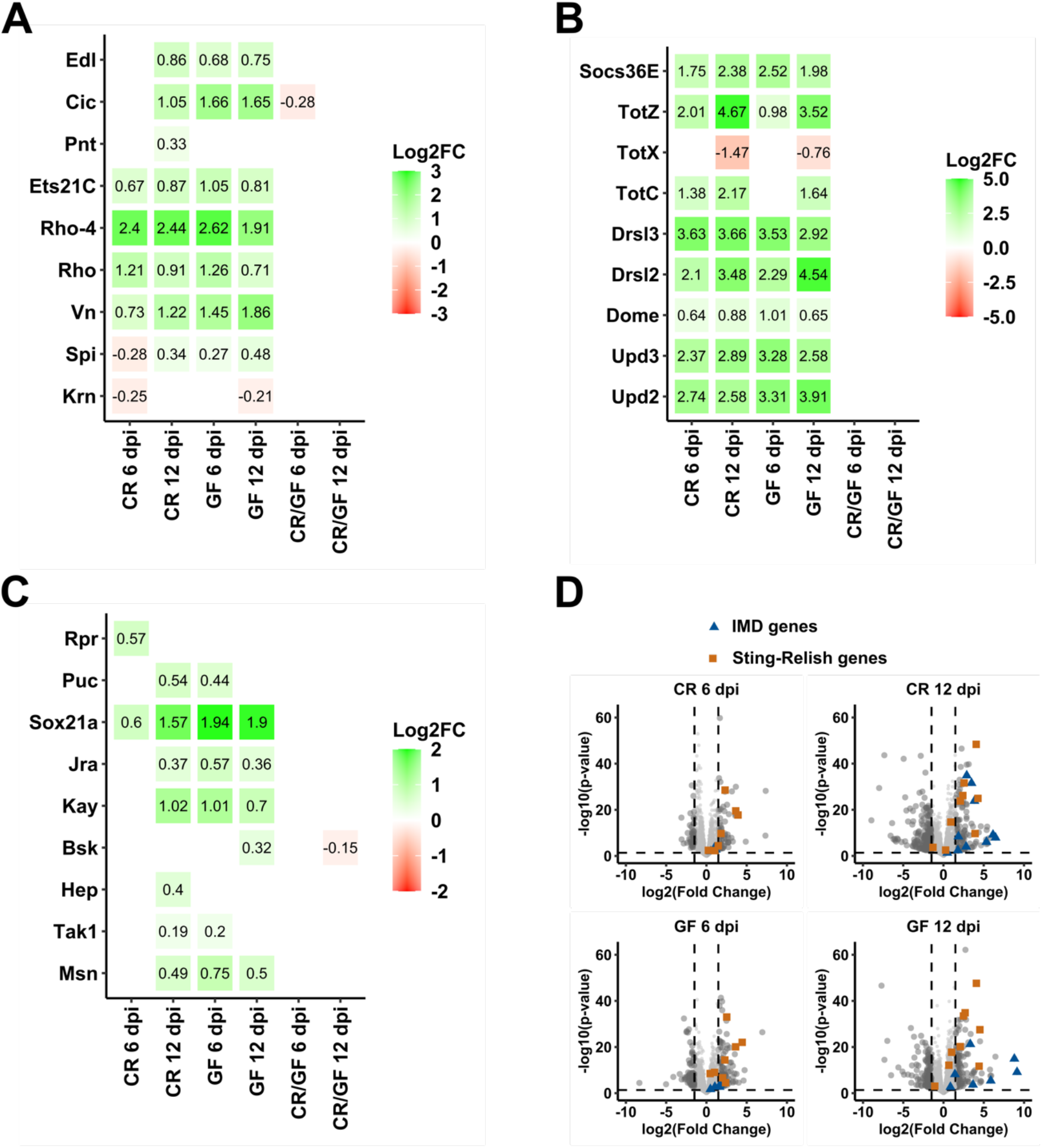
DAV infection upregulates immune, stress, and epithelial repair pathways in the intestine. (A-C) Expression of select genes of the EGFR (A), JAK-STAT (B), or JNK (C) pathways. Text and color indicate the log2 fold change (Log2FC) of expression in DAV-infected midguts/mock-infected midguts and in DAV-infected CR/GF midguts. Only genes with adjusted p-value < 0.05 are shown. (D) Expression of all genes. Select genes regulated by Imd-Relish or Sting-Relish signaling are highlighted. Expression in DAV-infected midguts/mock-infected midguts is shown. Horizontal dashed lines, adjusted p-value = 0.05; vertical dashed lines, Log2FC = 1.5. See also Figure S3. For complete data, see Data S1.

Besides EGFR and JAK-STAT signaling, the JNK pathway is another stress response pathway active in ECs and progenitors that serves distinct roles in each cell type to regulate ISC proliferation. JNK signaling is induced in ECs damaged by injury or pathogenic bacterial infection, where it induces apoptosis as well as release of EGFR ligands and Upd3^47^. These secreted molecules subsequently promote compensatory ISC proliferation by activating EGFR signaling in adjacent ISCs and JAK-STAT signaling in nearby visceral muscle cells, respectively^16,45,47,48^. Recent studies also demonstrated that progenitor-autonomous JNK activity can trigger ISC proliferation^47,49^. We found that the genes encoding the JNK pathway transcription factors Ets21C and Sox21a were upregulated in the midgut by DAV infection (Figures 3A and 3C). Additionally, DAV-infection induced modest and timepoint/condition-dependent upregulation of several other genes involved in the JNK signaling cascade (Figure 3C). Thus, our results suggest that DAV infection may induce classical midgut epithelial repair mechanisms by activating the EGFR, JAK-STAT, and JNK pathways.

We next investigated the potential immune mechanisms activated in the midgut by DAV infection. While RNA-interference is considered the primary antiviral response in arthropods^50^, inducible transcriptional responses are proposed to restrict viral replication in several insect species^30,51–61^. In *D. melanogaster,* the inducible immune response is classically defined by the Toll and Immune deficiency (IMD) pathways, both of which involve conserved nuclear factor-kB (NF-kB) signaling cascades that regulate antimicrobial effector expression^62^. Interestingly, several studies have recently uncovered roles for the IMD pathway in regulating intestinal homeostasis, suggesting that inducible immune responses may have unappreciated roles in regulating epithelial turnover^63–65^. The *D. melanogaster* genome encodes three NF-kB transcription factors: Relish, Dorsal, and Dorsal-related immunity factor (Dif). Transcriptional targets of Dorsal and Dif primarily include antimicrobial peptides (AMPs) induced in response to Toll pathway activation by fungi or gram-positive bacteria^66,67^, while Relish regulates a distinct set of AMPs in response to IMD pathway activation by gram-negative bacteria^68^. Genetic evidence implicates the Toll and IMD pathways in antiviral responses in *D. melanogaster*^30,51,52,59,61^, but it is unclear how Toll and IMD are activated by and restrict viral infection. We found that IMD pathway transcriptional targets were among the most upregulated genes in DAV-infected midguts at 12 dpi in both CR and GF flies (Figure 3D). A subset of these IMD-responsive genes was expressed significantly higher in DAV-infected midguts from CR flies compared to DAV-infected midguts from GF flies (Figure S3B), suggesting that microbiota input may have an additive effect on IMD activation during DAV infection. Toll pathway transcriptional targets were only modestly and inconsistently upregulated by DAV infection (Figure S3C).

Recent studies demonstrated that Relish can be activated through a second signaling cascade requiring the IMD pathway kinase, I-kappaN kinase β (IKKβ), but not the entire IMD pathway^69–72^. This novel pathway is induced during viral infection by two homologs of the mammalian cytoplasmic DNA sensor, cyclic GMP-AMP synthase (cGAS), termed cGAS-like receptor 1 and 2 (cGLR1 & cGLR2)^70,71^. Activated cGLR1 and cGLR2 produce cyclic dinucleotides (CDNs) that activate Sting (a homolog of the human protein, stimulator of interferon genes (STING)), which acts upstream from IKKβ to activate Relish^70–72^. This cascade, referred to here as Sting-Relish signaling, regulates transcription of genes distinct from those regulated by IMD signaling^69^. While the antiviral mechanisms of Sting-Relish signaling are unknown, this pathway is required for resistance to systemic infections with several RNA and DNA viruses and Sting-Relish signaling in ECs is antiviral against enteric DCV and Sindbis virus infections^69,72,73^. Several targets of Sting-Relish signaling were strongly upregulated by DAV infection in midguts from CR and GF flies at both 6 and 12 dpi (Figures 3D and S3D). Unlike IMD-responsive genes, there were no significant differences in the expression of genes regulated by Sting-Relish signaling in DAV-infected midguts from CR flies compared to DAV-infected midguts from GF flies (Figure S3D).

Together our RNA-seq data suggest that the midgut transcriptional response to DAV infection shares many similarities with the response to pathogenic bacterial infection, including activation of the EGFR, JAK-STAT, JNK, and IMD pathways. In line with our finding that DAV infection induces ISC proliferation in GF flies (Figure 2B), our data indicate that the microbiota does not significantly influence DAV-induced expression of EGFR, JAK-STAT, or JNK pathways genes (Figure 3). We found that DAV infection induces a strong NF-kB-dependent transcriptional response, with significant upregulation of both IMD- and Sting-Relish-responsive genes (Figures 3D and S3). Given the known roles of the EGFR, JAK-STAT, and JNK pathways in modulating epithelial turnover along with the emerging roles of the IMD pathway in regulating intestinal homeostasis, we decided to use these data as starting points to mechanistically investigate how DAV infection induces ISC proliferation.

### DAV-induced ISC proliferation requires EGFR but not JAK-STAT signaling

Our transcriptome sequencing results indicated that DAV infection upregulates canonical epithelial repair pathways in the intestine (Figure 3). We thus examined whether the mechanism of DAV-induced ISC proliferation is consistent with other stress-induced proliferative responses. We first depleted EGFR or JAK-STAT signaling in adult progenitors using *esg^ts^* to express RNAi constructs targeting *egfr*, the gene encoding the receptor of the EGFR pathway, or *Stat92e*, the gene encoding the sole transcription factor of the JAK-STAT pathway. Progenitor-specific depletion of EGFR signaling, but not JAK-STAT signaling, blocked both DAV-induced ISC proliferation and accumulation of Esg+ cells at 8 dpi without impacting viral RNA levels (Figures 4A-4C, S4A, and S4B). Depleting JAK-STAT signaling in progenitors reduced DAV-induced ISC proliferation compared to controls (Figure 4A), but the difference was not statistically significant. Because paracrine JAK-STAT signaling can regulate ISC proliferation non-cell-autonomously^41,42,45^, we ubiquitously expressed *Stat92e* RNAi using an *Actin-Gal4* driver (*Act-Gal4*). Ubiquitous depletion of JAK-STAT signaling did not significantly impact DAV-induced ISC proliferation or DAV RNA levels (Figures 4D and S4C). Moreover, *Upd3* mutants exhibited similar levels of DAV-induced ISC proliferation and DAV RNA as their isogenic wild-type counterparts (Figures S4D and S4E). Together these results demonstrate that cell-autonomous EGFR signaling is required for DAV-induced ISC proliferation and that JAK-STAT signaling does not play role in the proliferative response to DAV infection.

**Figure 4.**
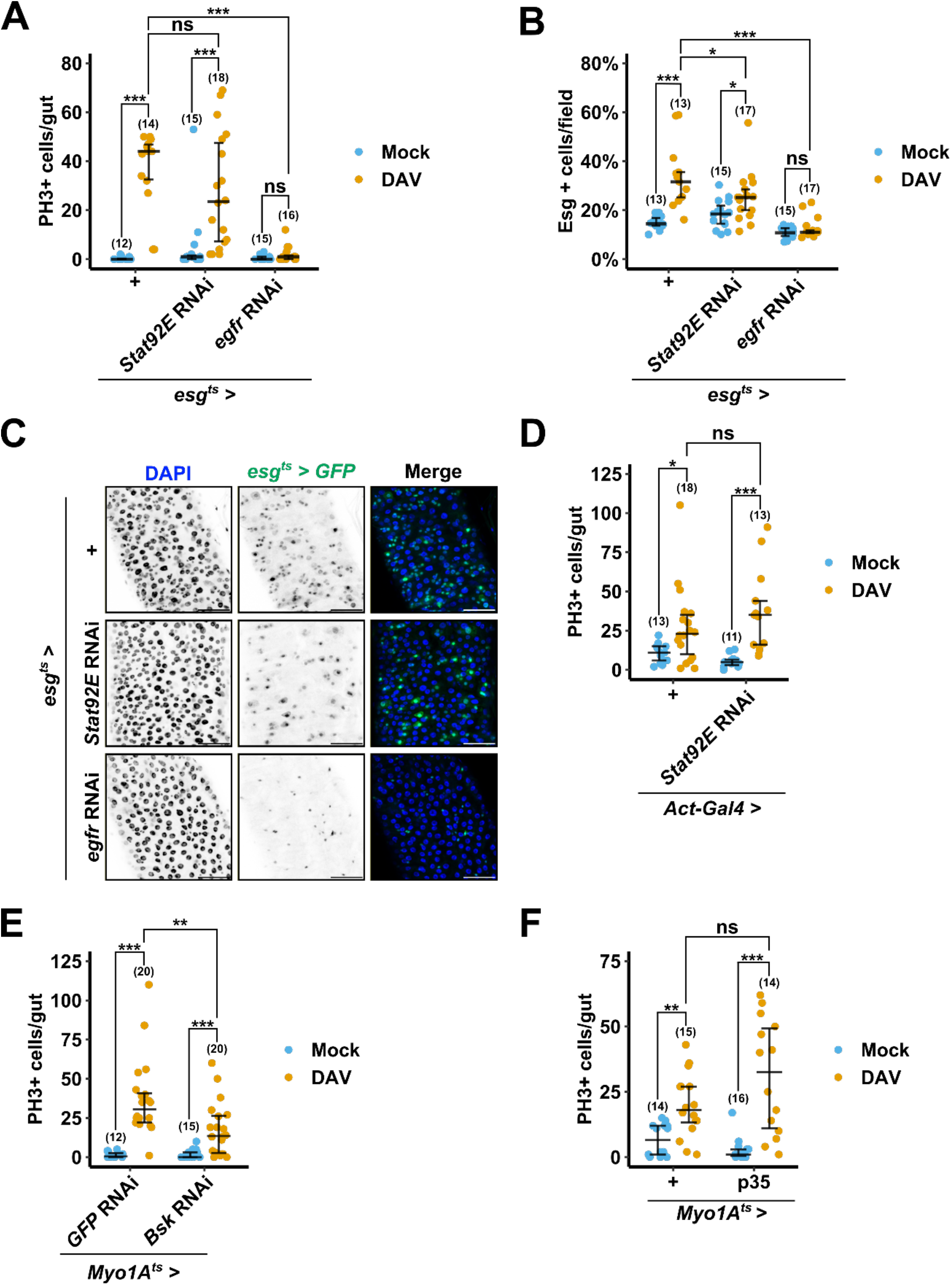
EGFR and JNK signaling, but not JAK-STAT signaling, play roles in DAV-induced ISC proliferation. (A) Quantification of PH3+ cells at 8 dpi in midguts from mock-infected or DAV-infected flies of the indicated genotypes. (B) Quantification of Esg+ cells in R4 midgut regions at 8 dpi from mock-infected or DAV- infected flies of the indicated genotypes. (C) Representative images of Esg+ cells at 8 dpi in R4 midgut regions from mock-infected or DAV-infected flies of the indicated genotypes. Scale bars: 50 μm. (D-F) Quantification of PH3+ cells at 8 dpi in midguts from mock-infected or DAV-infected flies of the indicated genotypes. Error bars indicate median with 1^st^ and 3^rd^ quartiles. Results were compared with a two-tailed Mann-Whitney test (A and D-F) or a two-tailed T-test (B); ns = non-significant, *p < 0.05, **p < 0.01, ***p < 0.001. Numbers of biological replicates indicated in parentheses. See also Figure S4.

### JNK signaling in ECs regulates DAV-induced ISC proliferation in an apoptosis- and JAK-STAT-independent manner

The genes encoding the JNK pathway transcription factors, Ets21C and Sox21a, were significantly upregulated during DAV infection (Figures 3A and 3C). To test a potential role for EC JNK activity in regulating DAV-induced ISC proliferation, we depleted JNK signaling in ECs by expressing an RNAi construct targeting the gene encoding Bsk, a serine/threonine-protein kinase responsible for phosphorylating the transcription factor encoded by *Jra* as part of the JNK signaling cascade. Using the temperature sensitive, EC-specific *Myo1A-Gal4 tubGal80^ts^*system (referred to as *Myo1A^ts^*)^16^ to express *Bsk* RNAi in adult ECs, we found that inhibition of JNK signaling in ECs did not prevent DAV-induced ISC proliferation, but did significantly reduce the levels of DAV-induced ISC proliferation compared to controls without impacting DAV RNA levels (Figures 4E and S4F).

Stress-induced apoptosis is an evolutionarily conserved antiviral mechanism and JNK-dependent apoptosis can trigger compensatory ISC proliferation in the *D. melanogaster* midgut^48,74^. However, a role for apoptosis of intestinal cells in restricting enteric viral infections in *D. melanogaster* has not been explored. We overexpressed the potent caspase inhibitor, p35^75^, using *Myo1A^ts^* to determine if EC apoptosis plays a role in DAV-induced ISC proliferation or restricting DAV infection. The levels of ISC proliferation were not significantly different in DAV-infected flies overexpressing p35 in ECs compared to controls (Figure 4F), indicating that the regulatory role of JNK activity in DAV-induced ISC proliferation is independent from EC apoptosis. Arguing against an antiviral role for caspase-dependent EC apoptosis during DAV infection, overexpressing p35 in ECs did not impact the relative median survival of DAV-infected flies and reduced the levels of DAV RNA compared to controls (Figures S4G and S4H). Together our results indicate that JNK activity in ECs plays a role in regulating DAV-induced ISC proliferation, but that JAK-STAT signaling, Upd3, and caspase-dependent EC apoptosis are dispensable for the role of the JNK pathway in this context.

### Sting-Relish signaling is required for DAV-induced ISC proliferation

Our results suggested that DAV infection induces ISC proliferation through mechanisms that overlap with but are incompletely described by canonical proliferative stress responses. Since NF-kB signaling is strongly upregulated in DAV-infected midguts (Figures 3D, S3B, and S3D), we infected flies with mutations in *Relish* (*Relish^E20^*, referred to as *Relish (-/-)*^68^) or *Dif* (*Dif^1^*, referred to as *Dif (-/-)*^76^) to determine if the IMD or Toll pathways, respectively, play roles in DAV-induced ISC proliferation. Strikingly, the absence of Relish completely abrogated DAV-induced ISC proliferation at 8 dpi despite higher DAV RNA levels in *Relish* mutants compared to isogenic wild-type flies (*w^1118^*, referred to as *WT^iso:Relish^*)(Figures 5A and S5A). DAV-infected *Dif* mutants exhibited significantly higher ISC proliferation levels compared to mock-infected controls, but significantly less than those in DAV-infected isogenic wild-type flies (*w^1118^,* referred to as *WT^iso:Dif^*) and there were no significant differences in DAV RNA levels between *Dif* mutants and wild-type flies (Figures S5B and S5C). In agreement with a previous report^77^, *Relish* mutants had reduced lifespans compared to wild-type flies (Figure S5D). To rule out the possibility that DAV-infected *Relish* mutants with a more proliferative ISC phenotype were already dead by 8 dpi, we measured ISC proliferation at 4 dpi. The absence of Relish prevented DAV-induced ISC proliferation at this earlier timepoint prior to the onset of mortality (Figure S5E). These results suggest that NF-kB signaling plays a role in DAV induced ISC proliferation, with a particular requirement for Relish in the proliferative response to DAV.

**Figure 5.**
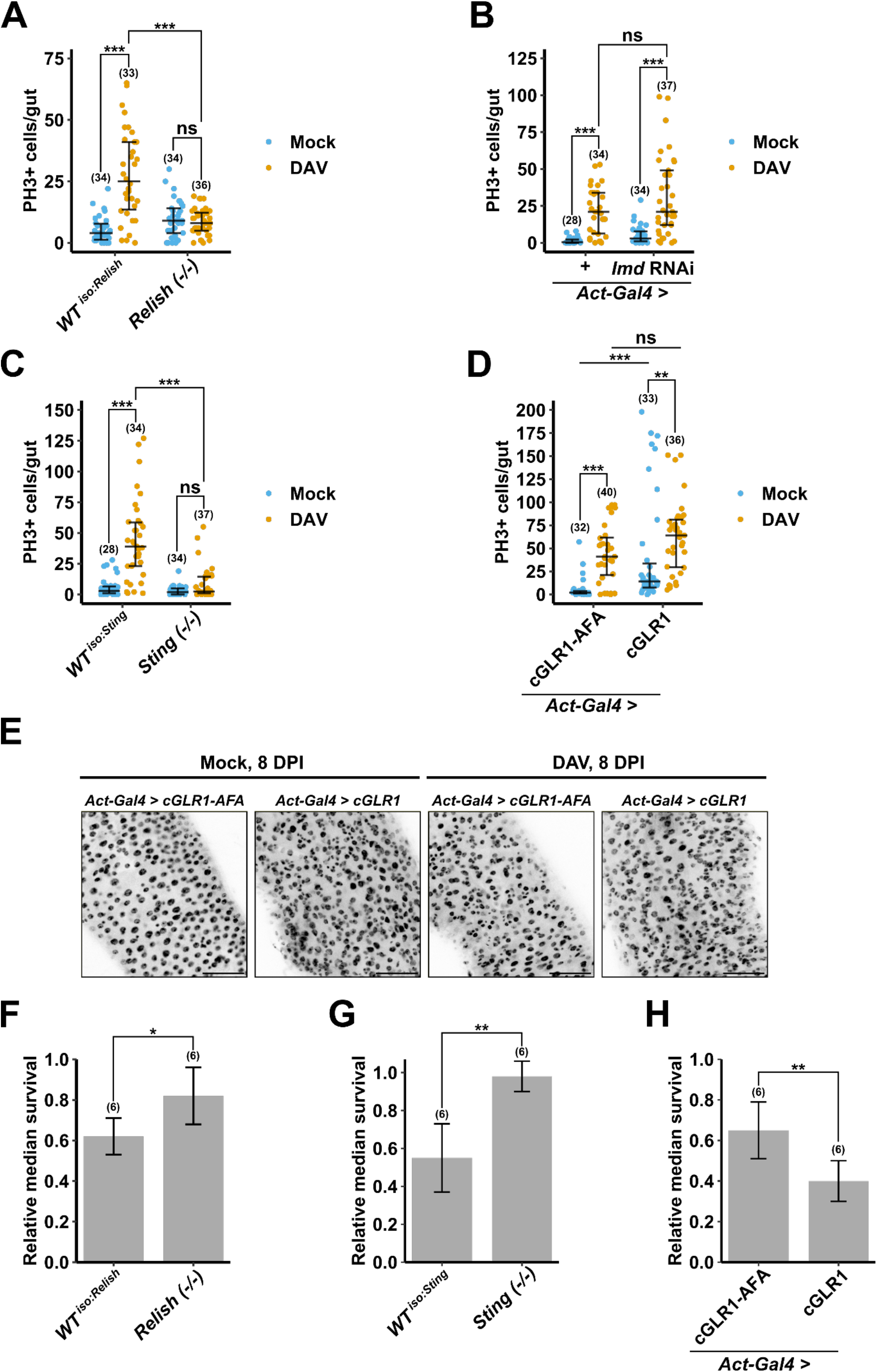
Sting-Relish signaling is required for DAV-induced ISC proliferation. (A-D) Quantification of PH3+ cells at 8 dpi in midguts from mock-infected or DAV-infected flies of the indicated genotypes. (E) Representative images of DAPI-stained R4 midgut regions at 8 dpi from mock-infected or DAV-infected flies of the indicated genotypes. Scale bars: 50 μm. (F-H) Relative median survival of DAV-infected flies of the indicated genotypes. Bar height indicates the average of biological replicates (n=15-20 flies/replicate). Error bars in (A-D) indicate median with 1^st^ and 3^rd^ quartiles. Error bars in (F-H) indicate S.D. Results were compared with a two-tailed Mann-Whitney test (A-D) or a two-tailed T- test (F-H); ns = non-significant, *p < 0.05, **p < 0.01, ***p < 0.001. Numbers of biological replicates indicated in parentheses. See also Figure S5.

Relish regulates both IMD- and Sting-dependent transcriptional responses. We thus infected *Sting* mutants (d*STING^Rxn^*, referred to as *Sting (-/-)*^69^) and flies with ubiquitously reduced expression of *Imd* (*Act-Gal4 Imd* RNAi) to investigate the relative contributions of Sting-Relish and IMD signaling, respectively, to Relish-dependent DAV-induced ISC proliferation. Infected *Imd*-knockdown flies exhibited similar levels of ISC proliferation and DAV RNA compared to infected controls at 8 dpi (Figures 5B and S5F). In contrast, DAV- infection did not induce significant levels of ISC proliferation in *Sting* mutants (Figure 5C). The lack of DAV-induced ISC proliferation in *Sting* mutants was not due to reduced DAV replication, as DAV RNA levels in *Sting* mutants compared to isogenic wild-type flies (*dSTING^control^*, referred to as *WT^iso:Sting^*) were not significantly different in whole flies or dissected midguts (Figures S5G and S6A).

Overexpression of cGLR1 or cGLR2 is sufficient to activate Sting-Relish signaling *in vivo*^71,78^. We thus compared ISC proliferation rates in flies ubiquitously overexpressing cGLR1 (*Act-Gal4* UAS-cGLR1^71^) to control flies overexpressing catalytically-inactive cGLR1 (*Act-Gal4* UAS-cGLR1-AFA^71^) to determine if Sting-Relish signaling is sufficient to induce ISC proliferation. Indeed, mock-infected flies overexpressing active cGLR1 had significantly greater ISC proliferation levels than mock-infected flies overexpressing catalytically inactive cGLR1 and exhibited epithelial irregularities resembling DAV-induced gut hypertrophy (Figures 5D and 5E). Notably, overexpressing cGLR1 had no impact on DAV RNA levels in infected flies (Figure S5H). Along with our finding that *Sting* knockout does not impact DAV RNA levels (Figures S5G and S6A), these data suggest that Sting-Relish signaling does not play an antiviral role during oral DAV infection. Together, our observations indicate that Sting-Relish signaling is required for DAV-induced ISC proliferation and is sufficient to promote ISC proliferation in the absence of infection.

Our results suggested that DAV infection may reduce lifespan by promoting sustained over-proliferation of ISCs, thus accelerating the onset of age-dependent intestinal pathology (Figures 2 and S1). If the reduced lifespan of DAV-infected flies is in fact driven by ISC proliferation, genetic interventions that preclude the proliferative response to DAV would be predicted to prolong lifespan. Since DAV-infection does not induce ISC proliferation in *Relish* or *Sting* mutants, we compared the relative median survival of DAV-infected *Relish* or *Sting* mutants to their wild-type counterparts. Indeed, the relative median survival of DAV-infected *Relish* or *Sting* mutants was significantly prolonged compared to wild-type flies, suggesting that Sting-Relish signaling may drive mortality during DAV infection (Figures 5F and 5G). In agreement with this possibility, ectopic activation of Sting-Relish signaling by overexpression of cGLR1 accelerated DAV-associated mortality and significantly reduced the lifespan of mock-infected flies (Figures 5H and S5I). Strikingly, the impact of cGLR1 overexpression on lifespan was equivalent to that of DAV infection (Figure S5I). Together our results indicate that NF-kB signaling plays a role in DAV-induced ISC proliferation and that Sting-Relish signaling is required for the proliferative response to DAV infection. Furthermore, our data support a role for virus-driven, Sting-dependent chronic NF-kB signaling in reducing lifespan by promoting sustained ISC proliferation, a phenotype that is recapitulated by activation of Sting-Relish signaling in the absence of infection.

### Loss of Relish or Sting diminishes DAV-induced upregulation of genes involved in the mitotic cell cycle and epithelial renewal in the intestine

We sequenced the transcriptomes of midguts dissected at 8 dpi from DAV- or mock-infected *Relish* or *Sting* mutants along with corresponding wild-type flies to identify differences in their transcriptional responses to DAV infection. We note that compared to midguts from wild-type flies, DAV RNA levels were significantly higher in midguts from *Relish* mutants, but not in midguts from *Sting* mutants (Figure S6A). As expected, loss of Relish or Sting reduced DAV-induced upregulation of transcriptional targets of the IMD-pathway and Sting-Relish signaling (Figures S6B and S6C). We identified 78 genes that were significantly upregulated by DAV infection in wild-type midguts at 8 dpi that were not significantly upregulated by DAV infection in midguts from *Relish* or *Sting* mutants (Figure 6A). In agreement with our ISC proliferation data in the mutants, functional enrichment analysis indicated that large portions of these genes belonged to the Gene Ontology (GO) terms, “multicellular organism development”, “mitotic cell cycle”, and “cell differentiation” (Figure 6A). Indeed, we observed broad upregulation of genes belonging to the GO term “mitotic cell cycle” in the intestines of DAV-infected wild-type flies, but not in DAV-infected *Relish* or *Sting* mutants (Figure 6B).

**Figure 6.**
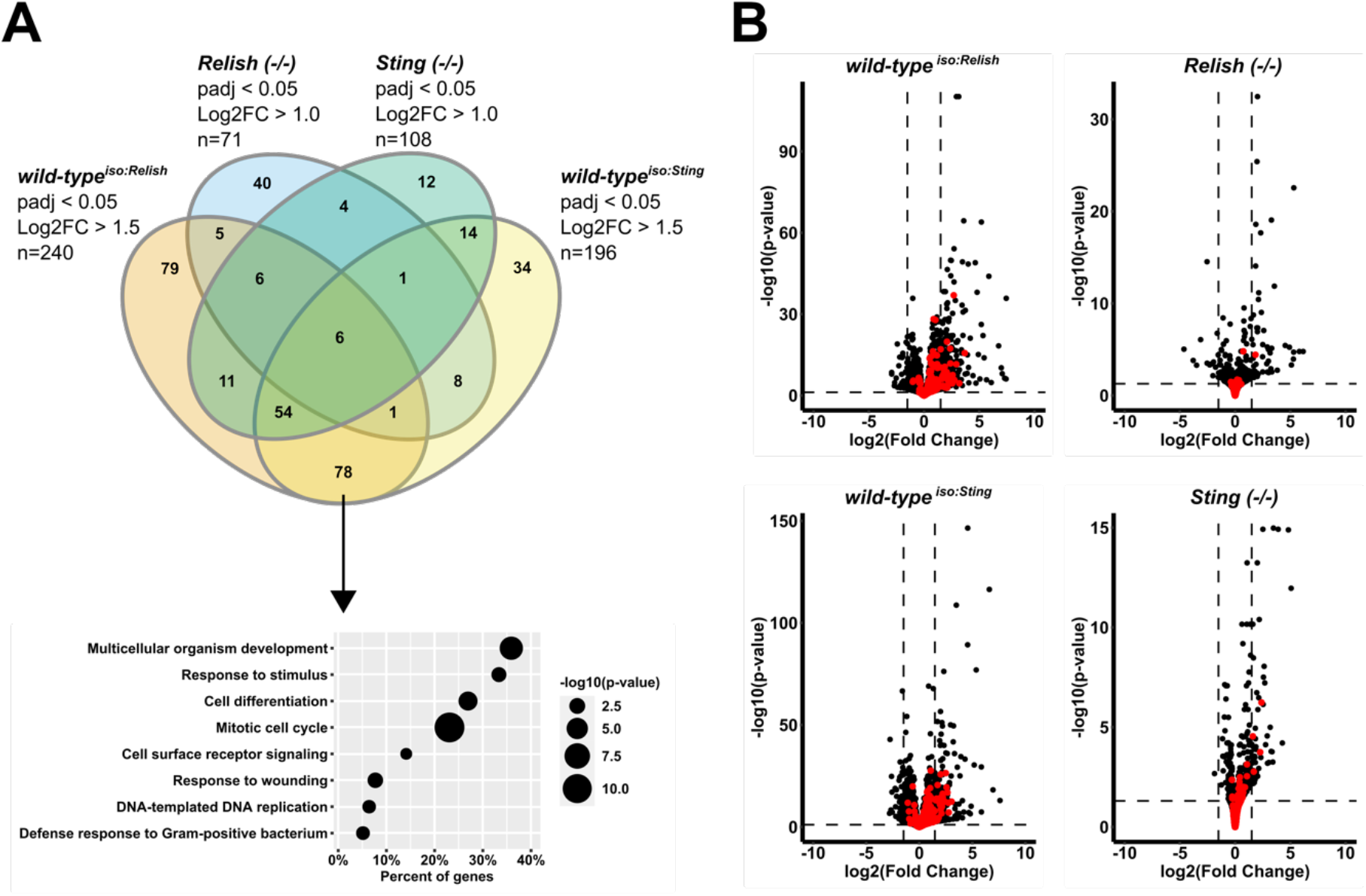
Loss of Relish or Sting diminishes DAV-induced upregulation of genes involved in the mitotic cell cycle and epithelial renewal in the intestine. (A) Upper: Overlap of genes differentially expressed in the midguts of flies with the indicated genotypes. Log2 fold change (Log2FC) is expression in DAV-infected/mock-infected conditions. Padj = adjusted p-value. Lower: select GO categories of genes upregulated in wild-type flies, but not in mutants. (B) Expression of all genes in the indicated genotypes. Genes in the GO category “mitotic cell cycle” are in red. Expression in DAV-infected midguts/mock-infected midguts is shown. Horizontal dashed lines, adjusted p-value = 0.05; Vertical dashed lines, Log2FC = 1.5. See also Figure S6. For complete data, see Data S2.

Our results indicated that progenitor-autonomous EGFR signaling and JNK signaling in ECs regulate ISC proliferation in response to DAV infection (Figures 4B and 4E). Compared to wild-type flies, the intestinal transcriptomes of DAV-infected *Relish* or *Sting* mutants were characterized by diminished upregulation of genes belonging to the EGFR and JNK pathways (Figures S6D and S6E). We also observed reduced upregulation of JAK-STAT pathway genes in midguts from DAV-infected *Relish* or *Sting* mutants compared to wild-type controls (Figure S6F). These findings are consistent with a previous study which found that ectopic activation of Sting-Relish signaling by injection of 2’3’-cGAMP induces upregulation of several key genes of the EGFR, JAK-STAT, and JNK pathways^72^. Together these results reveal that DAV infection induces global upregulation of genes involved in cell cycle progression, cellular differentiation, and regulation of epithelium renewal in the intestine. Loss of either Relish or Sting diminishes this transcriptional response to infection, supporting the possibility of a functional link between Sting-Relish signaling and canonical epithelial repair mechanisms during viral infection.

## DISCUSSION

In this study we leveraged the *D. melanogaster* model to elucidate the physiological consequences of enteric viral infection and investigate the host-virus interactions that influence infection outcomes. We found that orally acquired DAV persistently infects the adult midgut, induces sustained over-proliferation of ISCs, and accelerates age-associated intestinal pathology in a microbiota-independent manner (Figures 1, 2, and S1). We observed similar phenotypes in flies persistently infected with DCV, Nora virus, or Bloomfield virus, suggesting that modulation of intestinal physiology is a common feature of viral infections (Figure S1). Pathogenic bacterial infection induces a transient proliferative response in the *D. melanogaster* intestine, without which flies cannot repair the damaged epithelial layer and rapidly succumb to infection^15^. Blocking EC apoptosis did not influence the survival of infected flies and significantly reduced DAV RNA levels (Figure S4), arguing against an antiviral role for intestinal cellular turnover. In contrast, reducing the rate of ISC proliferation by ISC-specific overexpression of cell cycle inhibitors prolonged the survival of DAV-infected flies without impacting viral RNA levels (Figures 2 and S1), suggesting that elevated ISC proliferation is not an adaptive host response allowing flies to resist or tolerate DAV infection, but is instead a detrimental consequence of infection.

Is increased ISC proliferation beneficial for the viral infection cycle? One hypothesis is that loss of intestinal barrier function due to dysregulated cellular turnover may facilitate dissemination of infection beyond the midgut^79^. We observed that DAV RNA levels increased in extra-intestinal tissues prior to the onset of DAV replication in the gut and this pattern was not impacted by inhibiting ISC proliferation (Figures 1 and S1), suggesting that spread of DAV beyond the midgut does not depend on altered cellular turnover. Another possibility is that elevated ISC proliferation could modify epithelial composition in a way that promotes viral replication, for example, by increasing the density of susceptible cells and/or pro-viral host factors^80^. Similarly, increased ISC proliferation could conceivably facilitate changes to epithelial structure or function to promote shedding of infectious virus. Several lines of investigation arise from our data: Does DAV infection alter the relative proportions of differentiated intestinal epithelial cell types? Are metabolic or hormonal states modulated by viral infection? Are new cells produced by DAV-induced ISC proliferation retained? Does DAV infection influence non-apoptotic epithelial cell loss, such as engulfment^81^ and erebosis^82^. These questions should be addressed in future studies and may inform conclusions regarding the functional role of virus-driven modulation of cellular turnover.

RNA-seq of dissected midguts revealed that DAV infection induces upregulation of genes belonging to classical epithelial repair systems, including the EGFR, JAK-STAT, and JNK pathways (Figure 3). RNAi-based knockdown experiments indicated that DAV-induced ISC proliferation requires cell-autonomous EGFR signaling in progenitors and pointed towards an apoptosis-independent mitogenic role for JNK signaling in ECs during DAV infection (Figure 4). Unlike damage-induced proliferative responses, DAV-induced ISC proliferation did not depend on JAK-STAT signaling, suggesting that the role of JNK signaling in DAV- induced ISC proliferation is independent of the JAK-STAT pathway (Figure 4). Intriguingly, our results uncovered a requirement for Sting-dependent NF-kB signaling in the induction of ISC proliferation by DAV infection and constitutive activation of Sting-Relish signaling by systemic overexpression of cGLR1 was sufficient to induce ISC proliferation, promote gut hypertrophy, and reduce lifespan in the absence of infection (Figures 5 and S5)

Both IMD-Relish and Sting-Relish signaling are reported to play antiviral roles in *D. melanogaster*^52,59,61,69–73^. We observed increased DAV RNA levels in carcasses and midguts from *Relish* mutants, but not those from *Sting* mutants (Figures S5 and S6). Additionally, ubiquitous overexpression of cGLR1 did not impact DAV RNA levels or prolong survival of infected flies compared to controls (Figure S5). These results suggest that Relish is antiviral during enteric DAV infection, but that its antiviral role is independent from Sting-Relish signaling in this context. Critically, these results indicate that the requirement of Sting-Relish signaling for DAV-induced ISC proliferation is not an indirect result of reduced DAV RNA levels in the mutants. Supporting our ISC proliferation data, the midguts of DAV-infected wild-type flies exhibited broad upregulation of genes involved in the mitotic cell cycle and cell differentiation, a response that was not seen in *Relish* or *Sting* mutants (Figure 6). The midguts of DAV-infected *Relish* or *Sting* mutants also showed diminished upregulation of genes belonging to the EGFR, JAK-STAT, and JNK pathways compared to DAV-infected wild-type flies (Figure 6). Together our data suggest that Sting-Relish signaling may act either upstream from or in concert with classical epithelial repair pathways to promote ISC proliferation.

Although Relish is expressed in all intestinal cell types^83^, its function is best understood in ECs, where IMD-dependent Relish activation triggers production of AMPs^84^. IMD-Relish signaling can also synergize with the JNK pathway to control EC shedding during pathogenic bacterial infection^84^. While the roles of Relish in progenitors are less clear, a recent study found that Relish directly regulates expression of genes involved in cell cycle control and cellular differentiation in progenitors^85^. Accordingly, activation of IMD signaling in progenitors by commensal or pathogenic bacteria regulates ISC differentiation to establish the cellular composition of the epithelium^85^. Relish is not required for ISC proliferation in response to acute tissue damage, however several studies point towards a specific role for IMD-Relish signaling in promoting age-associated ISC proliferation^63–65^.

Sting-dependent Relish activation induces upregulation of genes distinct from those induced by the IMD-Relish pathway^69^, suggesting that Sting-Relish and IMD-Relish signaling may engage different factors to modify the regulatory activity of Relish and implying that the two signaling cascades have different functional outcomes. In mammals, cGAS-STING signaling splits into three branches downstream from STING to induce the type I interferon (IFN-1) response, activate NF-kB signaling, and trigger non-canonical autophagy^86^. Virus-induced production of IFN-1 is known to promote stem cell proliferation in the mouse intestine and NF-kB activity in mouse myeloid cells induces the production of tumor-promoting factors that stimulate intestinal epithelial cell proliferation in a paracrine manner^87,88^. Moreover, cGAS-STING-dependent IFN-β production in the ISC niche is required for compensatory proliferation of ISCs following acute radiation damage in mice^89^. Thus, investigation of inflammatory cGAS-STING and Sting-Relish signaling represent promising avenues to explore how host-microbe interactions modulate cellular turnover in epithelial tissues.

Persistent viral infections are common in arthropods and are known to be widespread in laboratory reared *D. melanogaster*, where Nora virus and DAV have reported prevalence of up to 52% and 18%, respectively^28^. Persistent viral infections are generally considered to pose no or only minor fitness costs for the host^27^, but there is relatively little published experimental data to support this claim and emerging studies demonstrate significant fitness impacts of persistent viral infection in *D. melanogaster*^90,91^. Moreover, re-analyses of published RNA-seq data from laboratory reared flies revealed previously undetected infections with several viruses including Nora virus, Thika virus, and DAV that induced significant changes in the host transcriptome^28,92^. Here we found that persistent infections with DAV, DCV, Nora virus, and Bloomfield virus all promote ISC proliferation and are associated with intestinal dysplasia (Figure S1). Together these results highlight that persistent viral infections can have significant phenotypic impacts and raise the possibility that undetected viral infections may have influenced previous experimental studies. In particular, we note that the microbiota-driven model of aging in *D. melanogaster* is based on experiments in which the commensal microbiota was ablated by dechorionation of embryos^15,18^. As this treatment also clears persistently infecting viruses^93^, one cannot rule out the possibility that viral infection may have contributed to the aging phenotypes previously ascribed solely to commensal dysbiosis. Given the high prevalence of persistent viral infection in laboratory flies and our observation that such infections can produce age-associated changes in the intestine, the potential contribution of viral infection to classical aging phenotypes should be studied in detail.

Our data reveal that persistent viral infection reduces lifespan by driving intestinal dysfunction in a manner involving sustained over-proliferation of ISCs. We propose that DAV infection accelerates aging by stimulating prolonged activation of inflammatory Sting-Relish signaling, resulting in dysregulation of classical epithelial repair systems and loss of intestinal homeostasis. Further studies are needed to elucidate the functional impacts of virus-driven modulation of epithelial turnover and to clarify the relationship between Sting-Relish signaling and epithelial repair pathways. Ultimately our results uncover wide-ranging impacts of viral infection on intestinal physiology and point towards novel host-virus interactions at the intersection of immune signaling, physiology, and aging.

## ACKNOWLEDGEMENTS

We thank J.L. Imler and P. Vale for fly stocks. This work was supported by funding from the French Government’s Investissement d’Avenir program, Laboratoire d’Excellence Integrative Biology of Emerging Infectious Diseases (grant ANR-10-LABX-62-IBEID), the Agence Nationale de la Recherche (grant ANR-23-CE15-0038-01, INFINITESIMAL), Fondation iXcore - iXlife - iXblue Pour La Recherche, and DIM One Health (Projet n° R17043DJ – Allocation n° RPH17043DJA) to M-C.S. This project has received funding from the European Union’s Horizon 2020 research and innovation program under the Marie Skłodowska*-*Curie grant agreement No 101024099 to J.N.

## AUTHOR CONTRIBUTIONS

J.N., V.M., X.G., A.J.B., and M.-C.S. designed the experiments; J.N. and H.B. performed the experiments; J.N., L.F., M.-C.S., and A.J.B. analyzed the data; J.N., A.J.B., and M.-C.S. wrote the paper. All authors reviewed and approved the final version of the manuscript.

## STAR METHODS

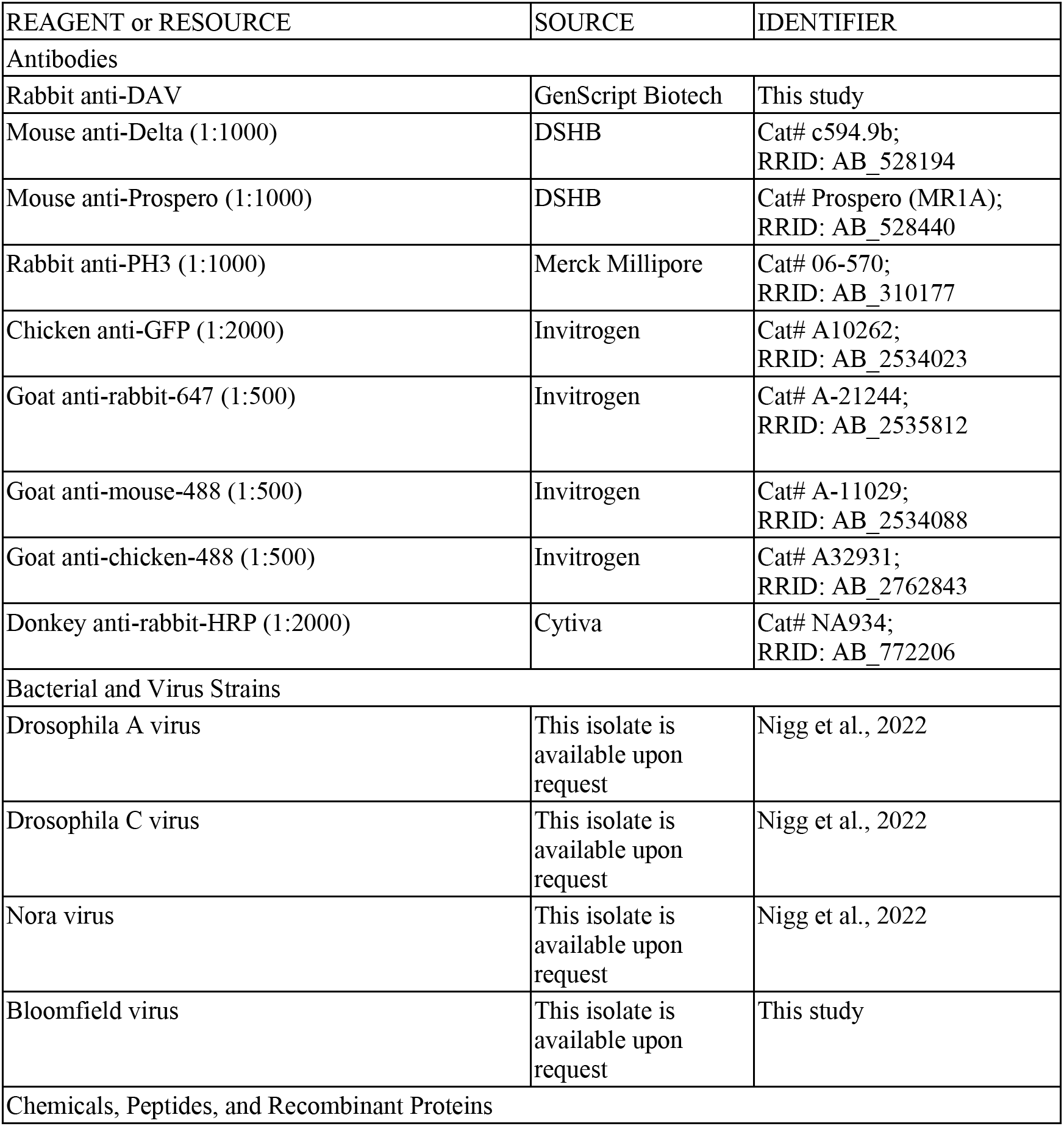
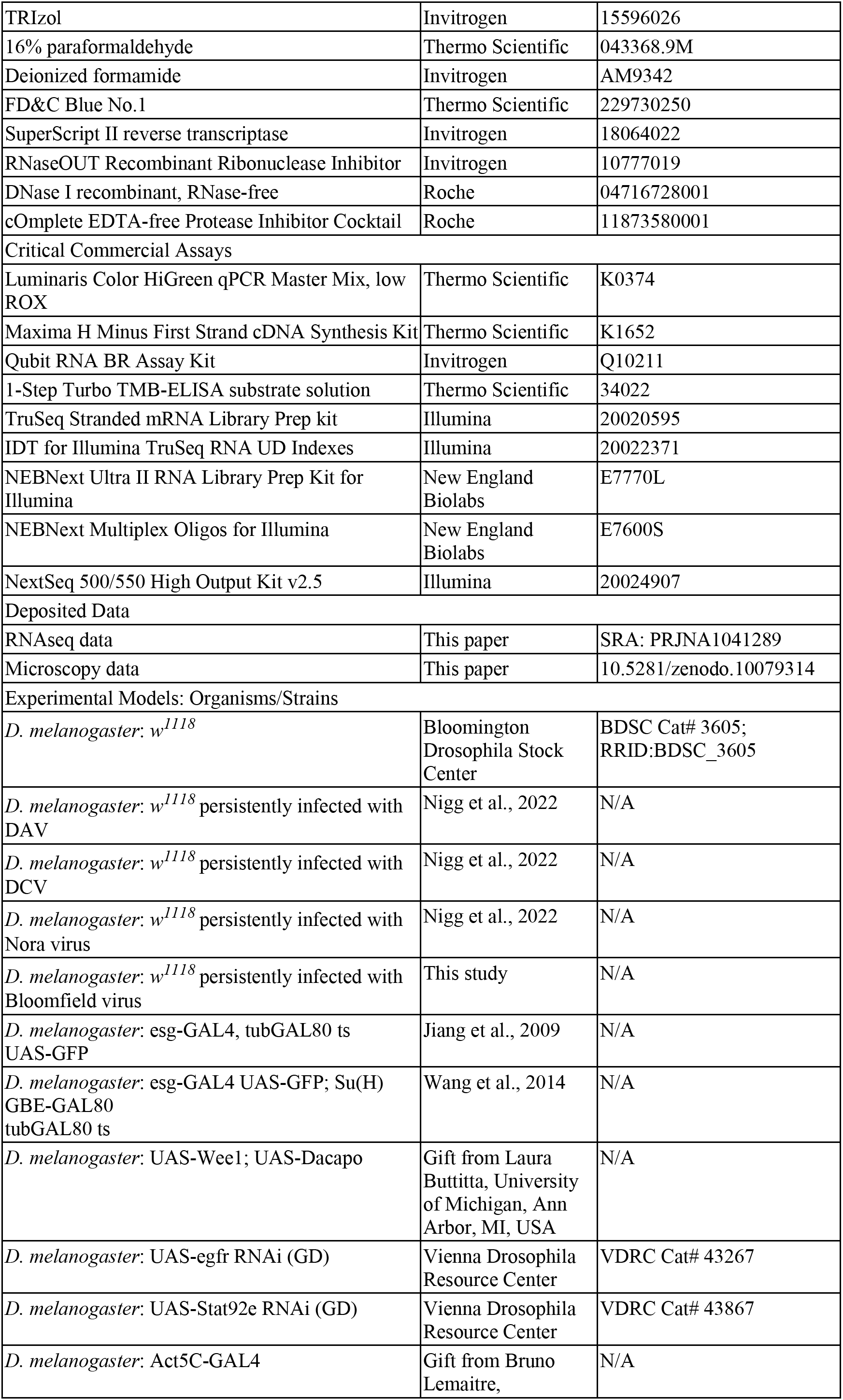
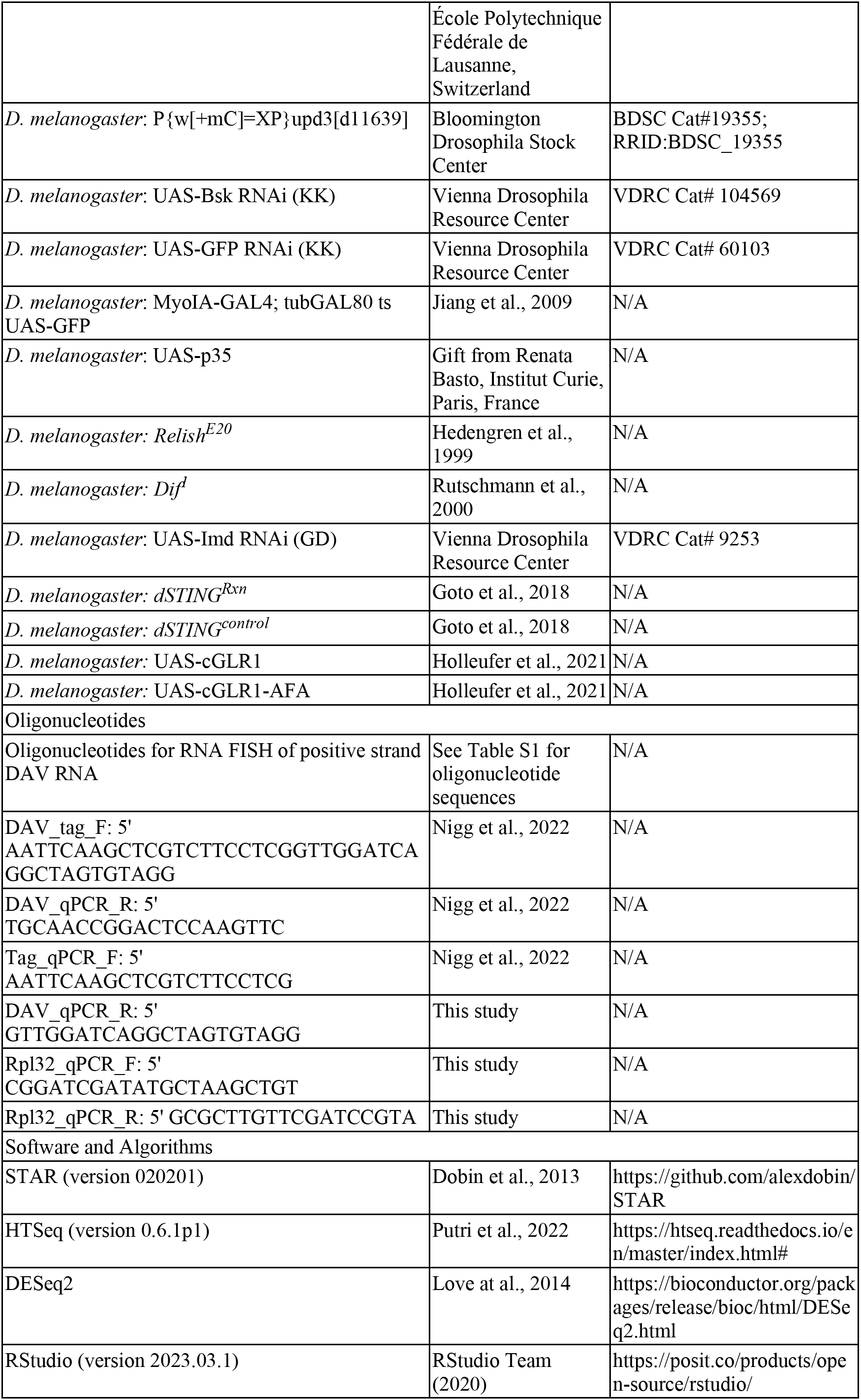
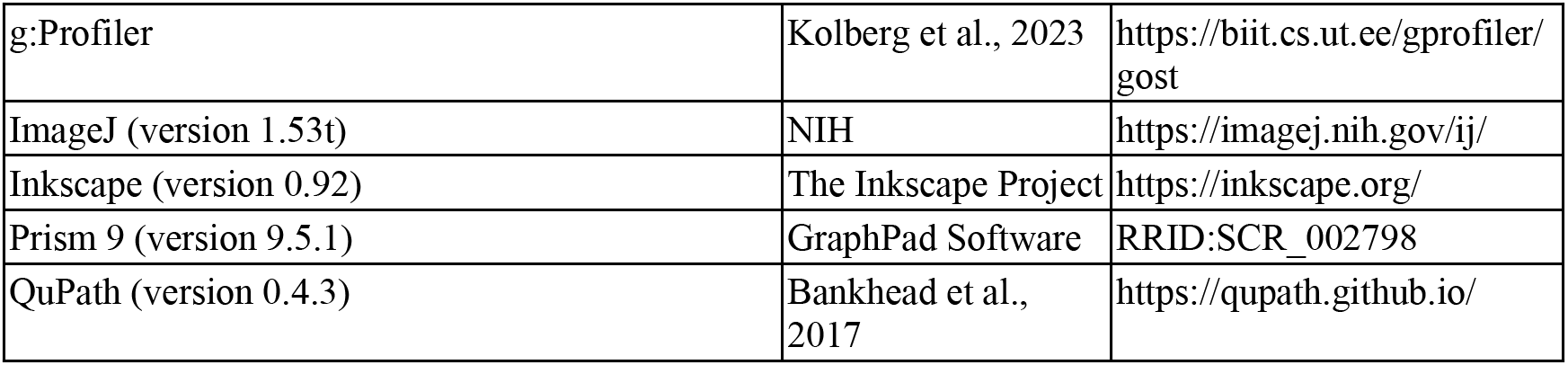
Key Resource Table.

### Resource availability

#### Lead contact

Further information and requests for resources and reagents should be directed to and will be fulfilled by the lead contact, Carla Saleh.

#### Materials availability

Materials generated in this study include persistently-infected fly stocks and an antibody against the DAV capsid protein. These are freely available without restriction from the lead contact upon request.

#### Data and code availability

- RNA-seq data have been deposited at SRA and are publicly available as of the date of publication. Accession numbers are listed in the key resource table.
- Microscopy data reported in this paper have been deposited at Zenodo and are publicly available as of the date of publication. DOIs are listed in the key resource table.
- This paper does not report original code.
- Any additional information required to reanalyze the data reported in this paper is available from the lead contact upon request.

### Experimental model and subject details

All fly stocks were maintained on a standard cornmeal diet (Bloomington) at 25 °C under a 12:12 hour light:dark cycle. All fly stocks were treated to clear potential persistent viral infections as previously described^93^. Briefly, eggs were treated with 50% bleach for 10 minutes, washed three times with distilled water, and transferred to fresh vials. All fly stocks were checked for the presence of *Wolbachia* infection as previously described and, when necessary, treated to clear Wolbachia infection^93^. This was accomplished by rearing the stocks on standard cornmeal diet containing 25 mg/mL tetracycline. Absence of *Wolbachia* infection was verified in the F3 of treated flies. Unless otherwise stated, *w^1118^* flies were used for all experiments. Fly stocks with mutations in *Relish* (*Relish^E20^*), *Dif* (*Dif^1^*), or *Upd3* were isogenized to the *w^1118^*line genetic background by backcrossing at least ten times to the *w^1118^*line as previously described^94^. Isogenized *Relish* and *Dif* mutants are the same stocks described by Mongelli et al.^94^. The isogenized *Upd3* mutant and the corresponding *w^1118^* wild-type stock were a gift from P. Vale (University of Edinburgh, Scotland). For Gal4-driven over-expression or RNAi experiments, virgin females containing the Gal4 driver were crossed with males containing the UAS-transgene and F1 adults (males and females together) were collected at 1-3 days of age. For experiments involving the *esg^ts^, esg^ts^; Su(H)-Gal80,* or *Myo1A^ts^* Gal4 driver lines, crosses were maintained at 18 °C. All other crosses were maintained at 25 °C. Gene expression or RNAi was induced by shifting flies to 29 °C 24 hours prior to infection. The effectiveness of gene silencing or overexpression was validated by RT-qPCR (data not shown). GF adult flies were generated by placing newly eclosed adults on standard cornmeal diet (Bloomington) containing 50 μg/ml ampicillin, 50 μg/ml kanamycin, and 10 μg/ml gentamycin. GF flies were maintained on the antibiotic-containing diet and flipped to fresh vials every two days. GF status was verified at 6 and 12 dpi by plating feces on LB agar plates. The fly stocks and crosses used for the experiments depicted in each figure are as follows.

- Figures 1, 2A, 2B, 2F, 3, S1A-F, S2: *w^1118^*
- Figures 2C-E: the F1 of *esg^ts^* females crossed with *w^1118^* males
- Figures 2G and H, S1G-I: the F1 of *esg^ts^; Su(H)-Gal80* females crossed with *w^1118^*or UAS-Wee1; UAS-Dacapo males
- Figures 4A-C, Figure S4A and B: the F1 of *esg^ts^* females crossed with *w^1118^*, UAS- Stat92e RNAi (VDRC Cat# 43867), or UAS-egfr RNAi (VDRC Cat# 43267) males
- Figure 4D and S4C: the F1 of *Act-Gal4* females crossed with UAS-Stat92e RNAi (VDRC Cat# 43867) males
- Figures 4E and S4F: the F1 of *Myo1A^ts^* females crossed with UAS-GFP RNAi (VDRC Cat# 60103) or UAS-Bsk RNAi (VDRC Cat# 104569) males
- Figures 4F and S4G and H: the F1 of *Myo1A^ts^* females crossed with *w^1118^* or UAS-p35 males
- Figures 5A and F and S5A, D, and E: *w^1118^* and isogenized *Relish^E20^*
- Figures 5B and S5F: the F1 of *Act-Gal4* females crossed with *w^1118^* or UAS-Imd RNAi (VDRC Cat# 9253) males
- Figures 5C and G and S5G: *dSTING^control^*and *dSTING^Rxn^* (gift from J.L. Imler, Université de Strasbourg, France)
- Figures 5D, E, and H and S5H and I: the F1 of *Act-Gal4* females crossed with UAS- cGLR1 or UAS-cGLR1-AFA males (gift from J.L. Imler, Université de Strasbourg, France)
- Figures 6 and S6: *w^1118^*, isogenized *Relish^E20^*, *dSTING^control^*, and *dSTING^Rxn^*
- Figures S5B and C: *w^1118^* and isogenized *Dif^1^*
- Figures S4D and E: *w^1118^*and isogenized P{w[+mC]=XP}upd3[d11639] (BDSC Cat#19355)

#### Generation of persistently infected flies

The *w^1118^* fly stocks persistently infected with DAV, DCV, or Nora virus have been described previously^33^. The *w^1118^* fly stock persistently infected with Bloomfield virus was generated as described previously^33^. Briefly, a Dipt-GFP reporter fly stock (BDSC Cat# 55709) was found to be contaminated with Bloomfield virus during routine screening. We homogenized the infected flies in PBS (5 μl/fly) and filtered the homogenate through a 0.22 μm filter. The homogenate was injected into 20 female and 10 male *w^1118^* flies (50 nl/fly) and the injected flies were placed in fresh vials containing standard cornmeal diet. After 3 days, the injected flies (F0) were removed and the F1 was reared in the same vial. The F1 flies were moved to a fresh vial 5 days after eclosion of the first adults. The F1 flies were removed from this vial after 9 days and F2 flies were moved to a new vial 5 days after eclosion of the first adults. The F2 was treated in the same manner and F3 adults were taken as the persistently infected stock. The presence of Bloomfield virus and the absence of other known *Drosophila-* infecting viruses was confirmed by total RNA sequencing.

### Method details

#### Viral infections

All infections were performed using groups of 20-40 mated adult female flies (3-5 days old). All infections were performed at 29 °C and infected flies were maintained at 29 °C unless otherwise stated. Flies were moved from their rearing temperature to 29 °C 24 hours prior to infection and starved for 5 hours prior to infection. Inoculation of standard cornmeal diet was performed by evenly coating the surface of the fly food with 100 µl of a DAV stock (1.3-2.0 x 10^3^ oral infectious dose 50% units/ml, see viral stock preparation and titration details below). Flies were placed on the contaminated food immediately following inoculation and were maintained on the contaminated food for 24 hours before being moved to fresh vials in groups of 20 flies/vial. The time at which flies were moved from the contaminated vials to fresh vials was considered as 0 dpi. Subsequently, the flies were flipped to fresh vials every two days. Mock-infections were performed in the same manner.

For experiments involving persistently-infected flies (as in Figures S1E and H), 10 males and 10 females of unknown age were collected from standard rearing vials for each persistently-infected stock or from standard rearing vials of uninfected *w^1118^*flies. These flies (referred to as the F0) were placed in fresh vials at 25 °C. 10 vials were setup in this manner for each fly stock and the F0 flies were removed after 10 days. The F1 flies (males and females together) were collected on the day of eclosion (referred to as 0 days-post eclosion) and moved to 29 °C. After 24 hours, the male F1 flies were removed and the female F1 flies were sorted into fresh vials at a density of 20 flies/vial. Subsequently, the flies were flipped to fresh vials every two days.

#### Antibody production

A polyclonal antibody against the DAV capsid protein (referred to as anti-DAV) was produced by GenScript Biotech (Piscataway, New Jersey, United States) using the PolyExpress Polyclonal Antibody Service. Briefly, New Zealand rabbits were immunized with the entire DAV capsid protein (GenBank accession no. YP_003038596) expressed in and purified from *E. coli* and anti-DAV was purified from the sera of immunized rabbit by affinity purification. The sensitivity and specificity of anti-DAV was verified by ELISA and western blot.

#### Viral stock preparation and titration

DAV stocks used for all experiments were prepared from *w^1118^* flies persistently infected with DAV. Flies of mixed ages and sex were collected from standard rearing vials and homogenized in 1x PBS (3 µl/fly). Homogenates were snap-frozen in a bath of dry ice/70% ethanol, stored overnight at -80 °C, thawed on ice, and clarified twice by centrifugation at 15,000 x g for 10 minutes at 4 °C. The clarified homogenate was filtered through a 0.22 µm filter and single use aliquots (100 µl) were snap-frozen in a bath of dry ice/70% ethanol and stored at -80 °C. A mock viral stock was prepared in the same manner using uninfected *w^1118^* flies.

DAV stocks were titered by 50% endpoint dilution via *in vivo* oral infection in adult flies to calculate an oral infectious dose 50% (OID50) for DAV stocks. The presence or absence of infection at 12 dpi was determined by ELISA in individual flies infected with a dilution series of DAV stock to calculate the OID50 according to the Reed and Muench method^95^. In detail, oral infections were performed as described above using ten-fold serially diluted DAV stocks ranging from undiluted to 10^-5^. Mock-infected flies served as a negative control. Individual flies (8 flies/dilution) were collected at 12 dpi and homogenized in 100 µl 1x PBS. 20 µl of each homogenate was mixed with 20 µl of lysis buffer (40 mM HEPES, pH 7.5, 2 mM DTT, 200 µM KCl, 10% glycerol, 0.1% NP-40, 1x cOmplete EDTA-free Protease Inhibitor Cocktail (Roche, 11873580001)) and incubated at room temperature for 15 minutes. 10 µl of the homogenate:lysis buffer mixture was added to 190 µl of 0.05 M carbonate-bicarbonate buffer, pH 9.6 in an Immuno 96-well flat bottom clear non-sterile plate, Nunc, MaxiSorp (Thermo Fisher, 442404) and incubated for 2 hours at room temperature. The plate was then washed three times with 200 µl/well of 1x PBS, 0.05% Tween-20 and incubated at room temperature for 2 hours with 200 µl/well of blocking buffer (1x PBS, 0.05% Tween-20, 5% non-fat dry milk). The plate was then washed three times with 200 µl/well of 1x PBS, 0.05% Tween-20 and incubated overnight at 4 °C with 100 µl/well of anti-DAV diluted 1:2000 in blocking buffer. The plate was then washed four times with 200 µl/well of 1x PBS, 0.05% Tween-20 and incubated at room temperature for 2 hours with 100 µl/well of donkey anti-rabbit IgG-HRP (Cytiva, NA934) diluted 1:2000 in blocking buffer. The plate was then washed four times with 200 µl/well of 1x PBS, 0.05% Tween-20 and incubated at room temperature for 2 hours with 100 µl/well of 1-Step Turbo TMB-ELISA substrate solution (Thermo Fisher, 34022). The reaction was stopped by addition of 100 µl/well of 2N HCl and absorbance at 450 nm was read on a Tecan Infinite M200 PRO plate reader. Infection status was determined for each fly based on A450 values and the threshold for infection was set as the average A450 for mock-infected flies plus ten times the standard deviation of A450 for the mock-infected flies. The OID50 of DAV stocks was calculated based on the infection prevalence for each dilution according to the Reed and Muench method^95^.

#### RNA extraction and RT-qPCR

Total RNA was extracted from whole flies, dissected midguts, or carcasses using TRIzol Reagent (Invitrogen, 15596026) according to the manufacturer’s instructions and RNA concentration was measured with the Qubit RNA BR Assay Kit (Invitrogen, Q10211). Carcasses included the entire fly body with the exception of the head and the alimentary canal.

DAV negative strand-specific RT-qPCR was performed as previously described^33^. Briefly RNA was reverse transcribed with DAV_tag_F (5’ AATTCAAGCTCGTCTTCCTCGGTTGGATCAGGCTAGTGTAGG) using SuperScript II Reverse Transcriptase (Invitrogen, 18064022) according to the manufacturer’s instructions with the following modifications: reverse transcription was performed at 50 °C for 30 minutes and reactions were heat inactivated for 15 minutes at 95 °C. cDNA was diluted 1:10 with distilled water and qPCR was performed in triplicate 10 µl reactions with the primers Tag_qPCR_F (5’ AATTCAAGCTCGTCTTCCTCG) and DAV_qPCR_R (5’ TGCAACCGGACTCCAAGTTC) using Luminaris Color HiGreen qPCR Master Mix, low ROX according to the manufacturer’s instructions (Thermo Scientific, K0374). qCPR was performed with a QuantStudio 7 Flex instrument (Applied Biosystems). The starting quantity of negative strand DAV RNA in each reverse transcription reaction was determined by absolute quantification by comparison to a standard curve generated by RT-qPCR of ten-fold serially diluted negative strand DAV RNA ranging from 10^2^ to 10^8^ copies/reaction. Samples below the limit of detection (10^3^ copies/reaction) were considered to have 0 negative strand copies. The number of negative strand copies per tissue was calculated from these data based on the RNA yield for each sample.

To calculate relative DAV RNA levels, RNA was treated DNase I recombinant, RNase-free (Roche, 04716728001) and reverse transcribed with Maxima H Minus First Strand cDNA Synthesis Kit (Thermo Scientific, K1652) according to the manufacturers’ instructions. The cDNA was diluted 1:10 with distilled water and amplified in triplicate 10 µl qPCR reactions for each sample and target using Luminaris Color HiGreen qPCR Master Mix, low ROX (Thermo Scientific, K0374) according to the manufacturer’s instructions (Thermo Scientific, K0374). DAV RNA was detected with the primers DAV_qPCR_F (5’ GTTGGATCAGGCTAGTGTAGG) and DAV_qPCR_R (5’ TGCAACCGGACTCCAAGTTC) and Rpl32 RNA was detected with the primers Rpl32_qPCR_F (5’ CGGATCGATATGCTAAGCTGT) and Rpl32_qPCR_R (5’ GCGCTTGTTCGATCCGTA). Relative DAV RNA levels were determined using the delta-delta Ct method^96^. DAV RNA levels in each sample were normalized to those of Rpl32 and are shown relative to the samples indicated in each figure legend.

#### Survival analysis

Survival analyses were conducted using biological replicates of 15-20 flies/replicate sorted at 1 dpi. All statistical comparisons involved flies maintained at identical initial population densities. Survival was monitored daily by counting the number of dead flies in each vial and flies were flipped to fresh vials every two days. For direct comparisons of survival curves (as in Figures 1B, S1D, S1G, S4F, and S4K), the survival data from all replicates of a given treatment and genotype were pooled and compared using a log-rank (Mantel-Cox) test. As expected, genotype and rearing condition (GF vs. CR) strongly influenced survival even under mock-infected conditions, confounding direct comparison of survival curves for DAV-infected flies of different genotypes or rearing conditions. Thus, we used the relative median survival to compare the influence of DAV infection on survival between different genotypes or rearing conditions while taking into account the different background survival rates of mock-infected flies (as in Figures 2H, 5F-H, S1B, and S3G). To calculate relative median survival, we first averaged the median survival (in days) of mock- or DAV-infected flies across biological replicates for a given genotype or treatment. We then calculated relative median survival as the average median survival across replicates of DAV-infected flies divided by the average median survival across replicates of mock-infected flies. For example, a relative median survival of 0.6 indicates that the average median survival of DAV-infected flies across biological replicates was 0.6 times that of mock-infected flies. The relative standard deviations of relative median survival values were calculated by propagation of error using the formula:

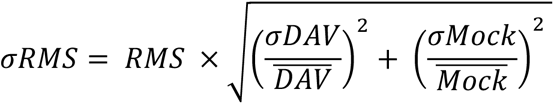

Where *σRMS* = standard deviation of the relative median survival, *RMS* = relative median survival, *σDAV* = standard deviation of the median survival for DAV-infected biological replicates, 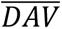 = average median survival of the DAV-infected biological replicates, *σMock* = standard deviation of the median survival for mock-infected biological replicates, and 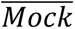 = average median survival of the mock-infected biological replicates. The values of *σRMS*, *RMS*, and sample size were used to determine the statistical significance of differences between relative median survival values using a two-tailed T-test.

#### Smurf assay

Smurf assays for measurement of intestinal barrier function were setup using biological replicates of 20 female *w^1118^* flies as described for survival analysis. Beginning from 7 dpi, flies were continuously maintained in vials in which 100 µl of a sterile solution of 32% FD&C blue dye #1 had been evenly spread on the surface of the diet. Following application of the blue dye solution, the vials were left to dry overnight at 25 °C before being used to house flies. Flies were flipped to fresh blue-dye treated vials every two days and “smurfness” was scored daily by examination of flies under a dissection microscope according to Rera et al.^97^. Dead flies were thoroughly washed with water to ensure that assessments of their blue coloration were based on internal blue dye rather than blue dye accumulated on external body surfaces. We calculated the median time (in days) until development of the smurf phenotype for each biological replicate and we evaluated the significance of differences in these median values between replicates of mock- and DAV-infected flies using a two-tailed T-test.

#### RNA FISH, immunofluorescence, and imaging

For detection of positive strand DAV RNA by RNA FISH, whole guts were dissected from mock- or DAV-infected flies in 1x PBS over a period of 20 minutes. Dissected guts were subsequently fixed in 4% paraformaldehyde, 0.3% Tween-20 for 20 minutes, washed twice for 2 minutes each time with 1x PBT (1x PBS, 0.1% Triton X-100), and then permeabilized by incubation in 1x PBS, 0.5% Triton X-100 for 20 minutes. Permeabilized tissue samples were washed for 2 minutes and then again for 10 minutes in fresh wash buffer (10% deionized formamide, 2x SSC) at room temperature. Finally, tissue samples were washed again with prewarmed wash buffer at 37 °C for 5 minutes and then incubated overnight with gentle shaking at 37 °C in prewarmed hybridization buffer containing 200 nM of 3’ ATTO 647-labeled DAV-specific oligonucleotides synthesized by DNA Script using a SYNTAX STX- 200 instrument^98^ (see Table S1 for oligonucleotide sequences, all oligonucleotides were mixed in equimolar amount). The hybridization buffer consisted of 10% deionized formamide, 5% dextran sulfate, 2x SSC, and 200 nM labeled oligonucleotides. The next day, tissue samples were washed for 2 minutes in fresh washer buffer and then rinsed three times for 10 minutes each time in 2x SSC. Subsequently, tissue samples were washed in 1x PBT for 2 minutes and then again for 10 minutes. DAPI was added to the second PBT wash at a final concentration of 1 µg/ml. Finally, tissue samples were mounted in 4% N-propyl-gallate, 80% glycerol.

For immunofluorescence of midgut tissues, midguts were dissected in 1x PBS over a period of 20 minutes and placed directly into 4% paraformaldehyde for fixation. Fixation continued for an additional 50 minutes following the end of the 20 minute dissection period. Fixed midguts were washed for 10 minutes three times (30 minutes total) in 1x PBT, incubated in 1x PBS, 50% glycerol for 30 minutes, and then equilibrated in 1x PBT for 30 minutes prior to incubation with primary antibodies diluted in 1x PBT. Primary antibody incubation occurred overnight at 4 °C. Midguts were then washed for 10 minutes three times before incubation with secondary antibodies diluted in 1x PBT. Secondary antibody incubation occurred for 3-5 hours at room temperature. Finally, midguts were washed for 10 minutes three times (the final wash contained 1 µg/ml DAPI) and mounted in 4% N-propyl-gallate, 80% glycerol. The following primary antibodies were used: anti-DAV (rabbit, 1:1000, generated in this study), anti-Delta (mouse, 1:1000, DSHB, c594.9b), anti-Prospero (mouse, 1:1000, DSHB, MR1A), anti-PH3 (rabbit, 1:1000, Merck Millipore, 06-570), anti-GFP (chicken, 1:2000, Invitrogen, A10262). The following secondary antibodies were used: anti-rabbit 647 (goat, 1:500, Invitrogen, A- 21244), anti-mouse-488 (goat, 1:500, Invitrogen, A-11029), anti-chicken-488 (goat, 1:500, Invitrogen, A32931). All imaging was performed on a Zeiss LSM 700 confocal microscope at the Institut Pasteur Unit of Technology and Service Photonic Bioimaging platform.

#### RNA-seq analysis

For the RNA-seq data depicted in Figures 3 and S2, CR or GF flies were infected as described above. Midguts were dissected at 6 and 12 dpi in 1x PBS. Midguts were dissected in 5 pools of 5 guts/pool, placed in 40 µl of ice-cold 1x PBS, and immediately transferred to dry ice upon dissection of each pool. RNA was extracted from midgut pools using 300 µl TRIzol reagent according to the manufacturer’s instructions. RNA-seq libraries were prepared with 150 ng RNA from 4 randomly selected pools/condition using a TruSeq Stranded mRNA Library Prep kit (Illumina, 20020595) with IDT for Illumina TruSeq RNA UD Indexes (Illumina, 20022371).

For the RNA-seq data depicted in Figures 6 and S5, flies were infected as described above. Midguts were dissected at 8 dpi. Individual midguts (12 midguts/condition) were dissected in 1x PBS, immediately transferred to 40 µl of ice-cold 1x PBS, and placed on dry-ice. RNA was extracted from individual midguts using 300 µl TRIzol reagent according to the manufacturer’s instructions. RNA concentrations were measured using the Qubit RNA BR Assay Kit (Invitrogen, Q10211). Within each condition, individual midguts were randomly assigned to 4 pools of 3 midguts/pool and 50 ng RNA from each individual midgut was combined to prepare the RNA pools. RNA-seq libraries were prepared from 150 ng of pooled RNA using an NEBNext Ultra II RNA Library Prep Kit for Illumina (New England Biolabs, E7770L) with NEBNext Multiplex Oligos for Illumina (Dual Index Primers Set 1) (New England Biolabs, E7600S). All sequencing was performed on an Illumina NextSeq 500 instrument using a NextSeq 500/550 High Output Kit v2.5 (75 cycles)(Illumina, 20024906).

Reads were mapped to the *D. melanogaster* genome (release dmel_r6.43) with STAR^99^ (version 020201). Feature counting was performed with HTSeq^100^ (version 0.6.1p1) using the default settings and differential gene expression analysis was performed with DESeq2^101^ in R Studio (version 2023.03.1). Log2 fold changes were shrunken using the ashr algorithm and only adjusted p-values were considered for analysis.

#### Gene Ontology enrichment analysis

Gene Ontology enrichment analysis was performed with the g:GOst function of g:Profiler^102^ using the default settings.

### Quantification and statistical analysis

All plots of data were prepared using ggplot2 in R Studio (version 2023.03.1). Confocal images were minimally processed in ImageJ (version 1.53t). Adjustments made to raw images included cropping, annotation, and adjustments to brightness and contrast applied across the entire image. Figures were assembled in Inkscape (version 0.92). Statistical analyses were performed using the GraphPad Prism software (version 9.5.1). Mitotic cells (PH3+ cells) were counted manually in the entire midgut and analyzed using two-tailed Mann-Whitney tests. For scoring of Esg+ cells, total nuclei and Esg+ nuclei were identified using QuPath^103^ (version 0.4.3) in unprocessed confocal images acquired at 25x from R2 and R4 midgut regions. Analysis regions were defined manually for each image to include the entire region of the midgut present in the field, but to exclude non-target tissues that were occasionally present in the field (malpighian tubules, non-target midgut regions present due to the orientation of the mounted tissue sample, and other midguts located nearby on the slide). Proportions of Esg+ cells were analyzed using two-tailed T-tests. Survival was analyzed by log-rank (Mantel-Cox) tests and two-tailed T-tests as described under survival analysis. Negative-strand DAV RNA levels and relative DAV RNA levels obtained by RT-qPCR were analyzed using two-tailed T-tests. Complete details regarding the statistical tests used, sample sizes, dispersion and precision measures, and definitions of significance are reported in the figure legends.

---

**Figure S1.**
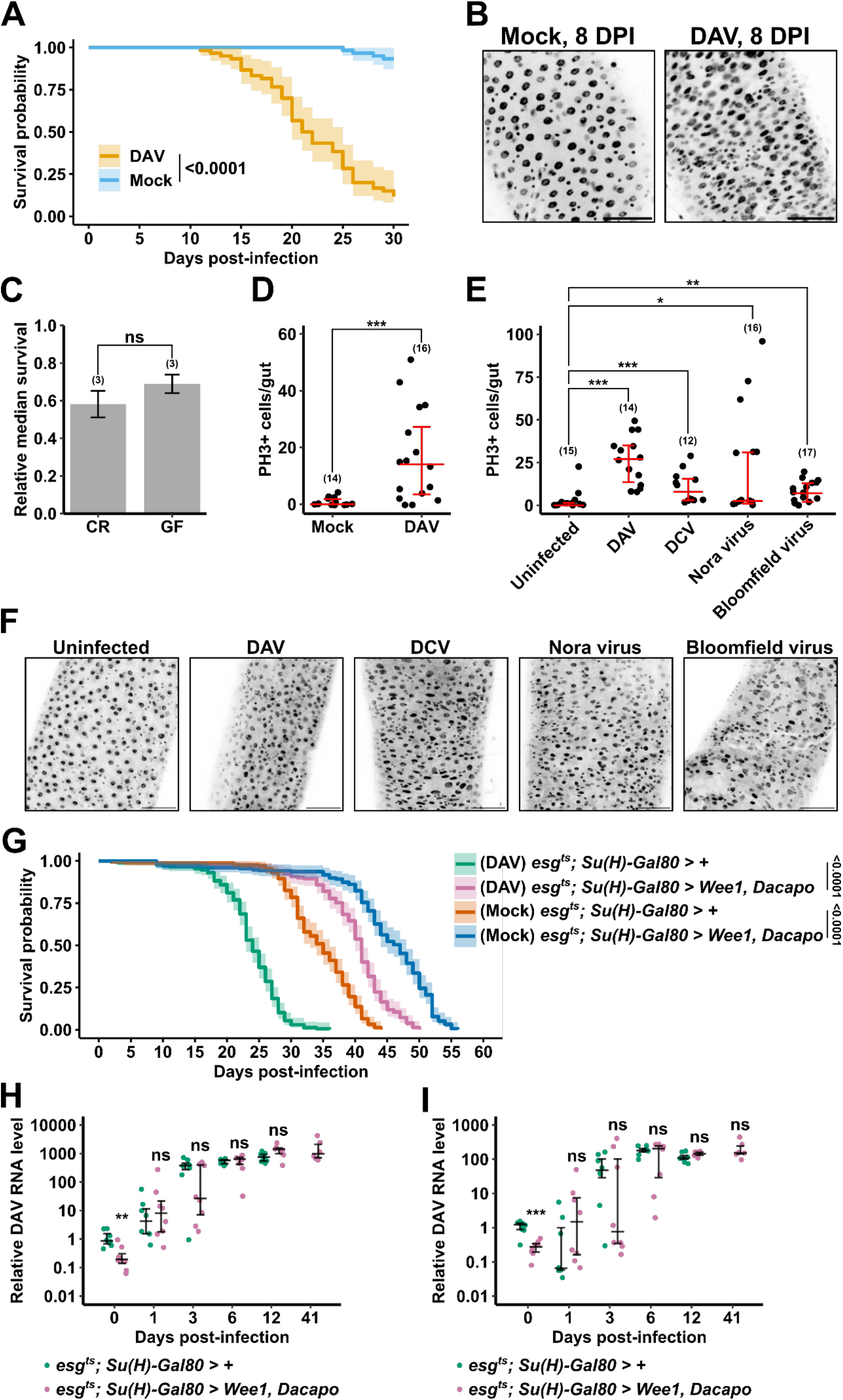
(A) Survival of mock-infected and DAV-infected flies maintained at 25 °C. Shaded regions: 95% confidence intervals. Three biological replicates (n=20 flies/replicate) were analyzed. The p-value from a log-rank (Mantel-Cox) test is shown. (B) Representative images of DAPI-stained R4 midgut regions at 8 dpi from mock-infected or DAV-infected flies maintained under GF conditions. Scale bars: 50 μm. (C) Relative median survival of DAV-infected flies maintained under CR or GF conditions. Bar height indicates the average of biological replicates (n=20 flies/replicate). (D) Quantification of PH3+ cells at 8 dpi in midguts from mock-infected or DAV-infected flies maintained at 25 °C. (E) Quantification of PH3+ cells at 20 days-post eclosion in midguts from uninfected *w^1118^* flies or isogenic *w^1118^* flies persistently infected with DAV, DCV, Nora virus, or Bloomfield virus. (F) Representative images of DAPI-stained R4 midgut regions at 20 days-post eclosion from uninfected *w^1118^* flies or isogenic *w^1118^* flies persistently infected with DAV, DCV, Nora virus, or Bloomfield virus. Scale bars: 50 μm. (G) Survival of mock-infected and DAV-infected flies of the indicated genotypes. Shaded regions: 95% confidence intervals. Nine biological replicates (n=20 flies/replicate) from three independent experiments were analyzed. The p-values from log-rank (Mantel-Cox) tests are shown. (H and I) Relative DAV RNA levels in carcasses or dissected midguts from DAV-infected flies of the indicated genotypes. DAV RNA levels are shown relative to the *esg^ts^; Su(H)- Gal80 > +* 0 dpi samples. n=12 samples/day for 0-12 dpi. n=6 samples for 41 dpi. Data at 41 dpi was not available for DAV-infected flies with the genotype *esg^ts^; Su(H)-Gal80 > +* because they had 100% mortality by 32 dpi. Error bars in (B) indicate S.D. Error bars in (C, E, G, and H) indicate median with 1^st^ and 3^rd^ quartiles. Results were compared with a two-tailed T-test (B, G, and H) or a two-tailed Mann-Whitney test (C and E); ns = non-significant, *p < 0.05, **p < 0.01, ***p < 0.001. Comparisons in (G and H) are between mock and DAV for each dpi. Numbers of biological replicates indicated in parentheses. See also Figures 1 and 2.

**Figure S2.**
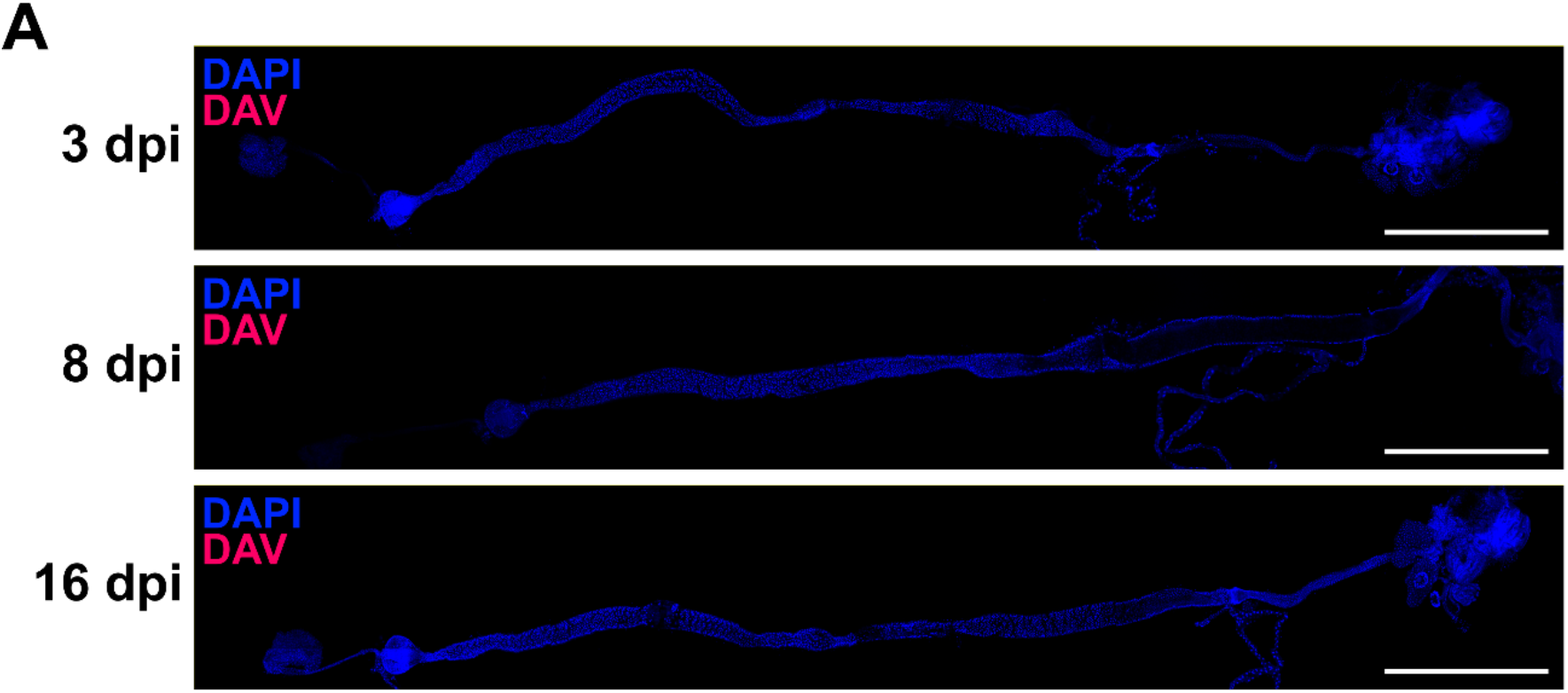
(A) RNA FISH of positive strand DAV RNA in guts from mock-infected flies. Scale bars: 1 mm. See also Figure 1.

**Figure S3.**
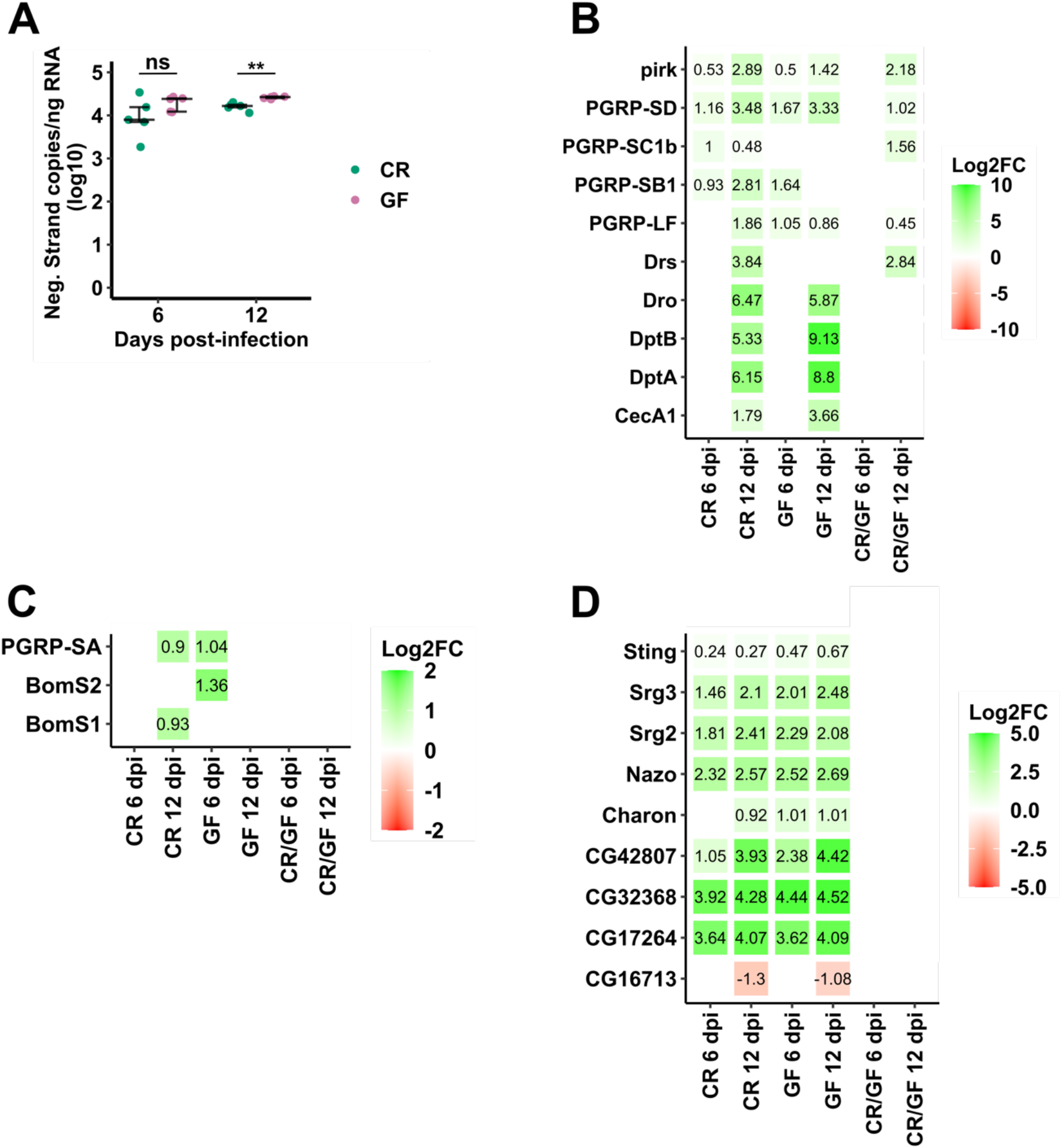
(A) Negative strand-specific RT-qPCR of DAV RNA in dissected midguts from DAV-infected flies maintained under CR or GF conditions. n=5 pools of 5 midguts/pool for each time and condition. Error bars indicate median with 1^st^ and 3^rd^ quartiles. Results were compared with a two-tailed t-test; ns = non-significant, **p < 0.01. (B-D) Expression of select genes regulated by the IMD pathway (A), Toll pathway (B), or Sting-Relish signaling (C). Text and color indicate the log2 fold change (Log2FC) of expression in DAV-infected midguts/mock-infected midguts and in DAV-infected CR/GF midguts. Only genes with adjusted p-value < 0.05 are shown. See also Figure 3.

**Figure S4.**
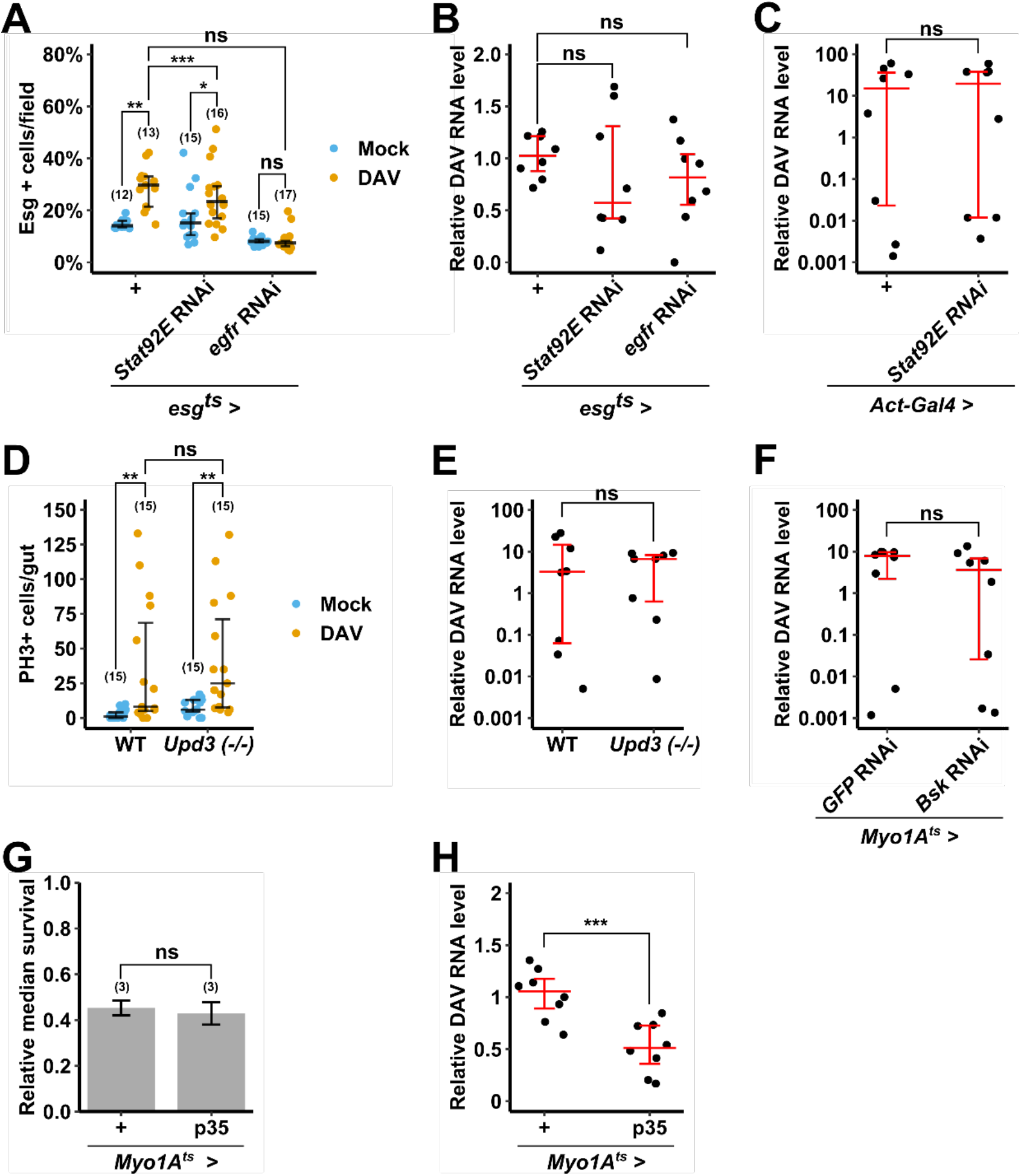
(A) Quantification of Esg+ cells in R2 midgut regions at 8 dpi from mock-infected or DAV-infected flies of the indicated genotypes. (B and C) Relative DAV RNA levels at 8 dpi in DAV-infected flies of the indicated genotypes. DAV RNA levels are shown relative to the *esg^ts^ > +* (B) or *Act-Gal4 > +* (C) samples. (D) Quantification of PH3+ cells at 8 dpi in midguts from mock-infected or DAV-infected flies of the indicated genotypes. (E and F) Relative DAV RNA levels at 8 dpi in DAV-infected flies of the indicated genotypes. DAV RNA levels are shown relative to the WT (E) or *Myo1A^ts^ > GFP* RNAi (F) samples. (G) Relative median survival of DAV-infected flies maintained of the indicated genotypes. Bar height indicates the average of biological replicates (n=20 flies/replicate). (H) Relative DAV RNA levels at 8 dpi in DAV-infected flies of the indicated genotypes. DAV RNA levels are shown relative to the *Act-Gal4 > +* samples. Error bars in (A-F and H) indicate median with 1^st^ and 3^rd^ quartiles. Error bars in (G) indicate S.D. Results were compared with a two-tailed Mann-Whitney test (A and D) or a two-tailed T-test (B, C, and E-H); ns = non-significant, *p < 0.05, **p < 0.01, ***p < 0.001. Numbers of biological replicates indicated in parentheses for (A, D, and G). n = 8 individuals flies for (B, C, E, F, and H). See also Figure 4.

**Figure S5.**
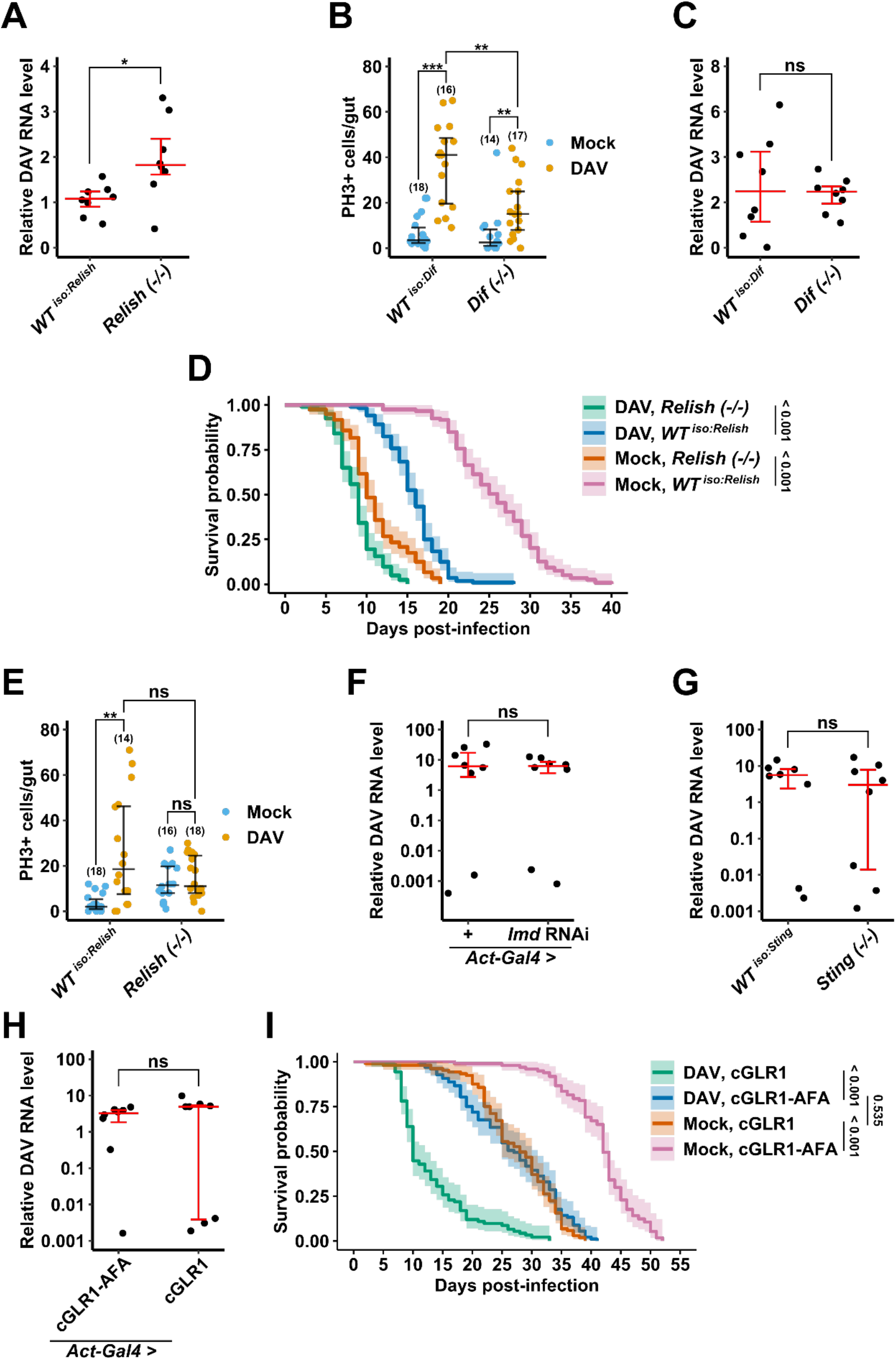
(A) Relative DAV RNA levels at 8 dpi in DAV-infected flies of the indicated genotypes. DAV RNA levels are shown relative to the *WT ^iso:Relish^* samples. (B) Quantification of PH3+ cells at 8 dpi in midguts from mock-infected or DAV-infected flies of the indicated genotypes. (C) Relative DAV RNA levels at 8 dpi in DAV-infected flies of the indicated genotypes. DAV RNA levels are shown relative to the *WT ^iso:Dif^* samples. (D) Survival of mock-infected and DAV-infected flies of the indicated genotypes. Shaded regions: 95% confidence intervals. Six biological replicates (n=20 flies/replicate) were analyzed. The p-values from log-rank (Mantel-Cox) tests are shown. (E) Quantification of PH3+ cells at 4 dpi in midguts from mock-infected or DAV-infected flies of the indicated genotypes. (F-H) Relative DAV RNA levels at 8 dpi in DAV-infected flies of the indicated genotypes. DAV RNA levels are shown relative to the *Act-Gal4 > +* (F), *WT ^iso:Sting^* (G), or *Act-Gal4* > cGLR1-AFA (H) samples. (I) Survival of mock-infected and DAV-infected flies of the indicated genotypes. Shaded regions: 95% confidence intervals. Six biological replicates (n=15-20 flies/replicate) were analyzed. The p-values from log-rank (Mantel-Cox) tests are shown. Error bars in indicate median with 1^st^ and 3^rd^ quartiles. Results were compared with a two-tailed T-test (A, C, and F-H) or a two-tailed Mann-Whitney test (B and E); ns = non-significant, *p < 0.05, **p < 0.01, ***p < 0.001. Numbers of biological replicates indicated in parentheses for (B and D). n = 8 individuals flies for (A, C, and F-H). See also Figure 5.

**Figure S6.**
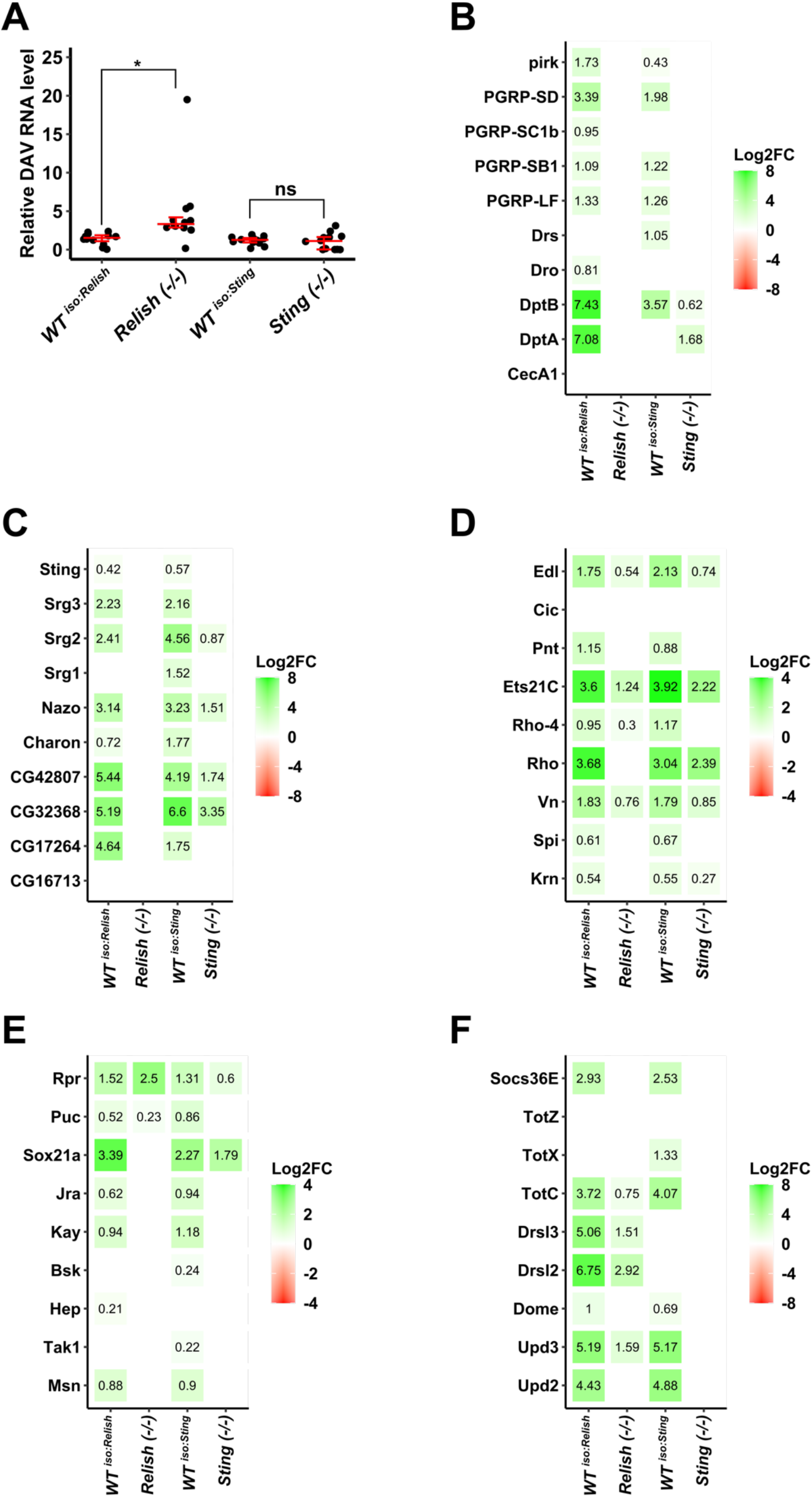
(A) Relative DAV RNA levels at 8 dpi in dissected midguts from DAV-infected flies of the indicated genotypes. n=12 individual midguts. Error bars indicate median with 1^st^ and 3^rd^ quartiles. Results were compared with a two-tailed t-test; ns = non-significant, *p < 0.05. (B-F) Expression of select genes regulated by the IMD pathway (B), Sting-Relish signaling (C), the EGFR pathway (D), the JNK pathway (E), or the JAK-STAT pathway (F). Text and color indicate the log2 fold change (Log2FC) of expression in DAV-infected midguts/mock-infected midguts from flies of the indicated genotypes. Only genes with adjusted p-value < 0.05 are shown. See also Figure 6.

**Table S1.**
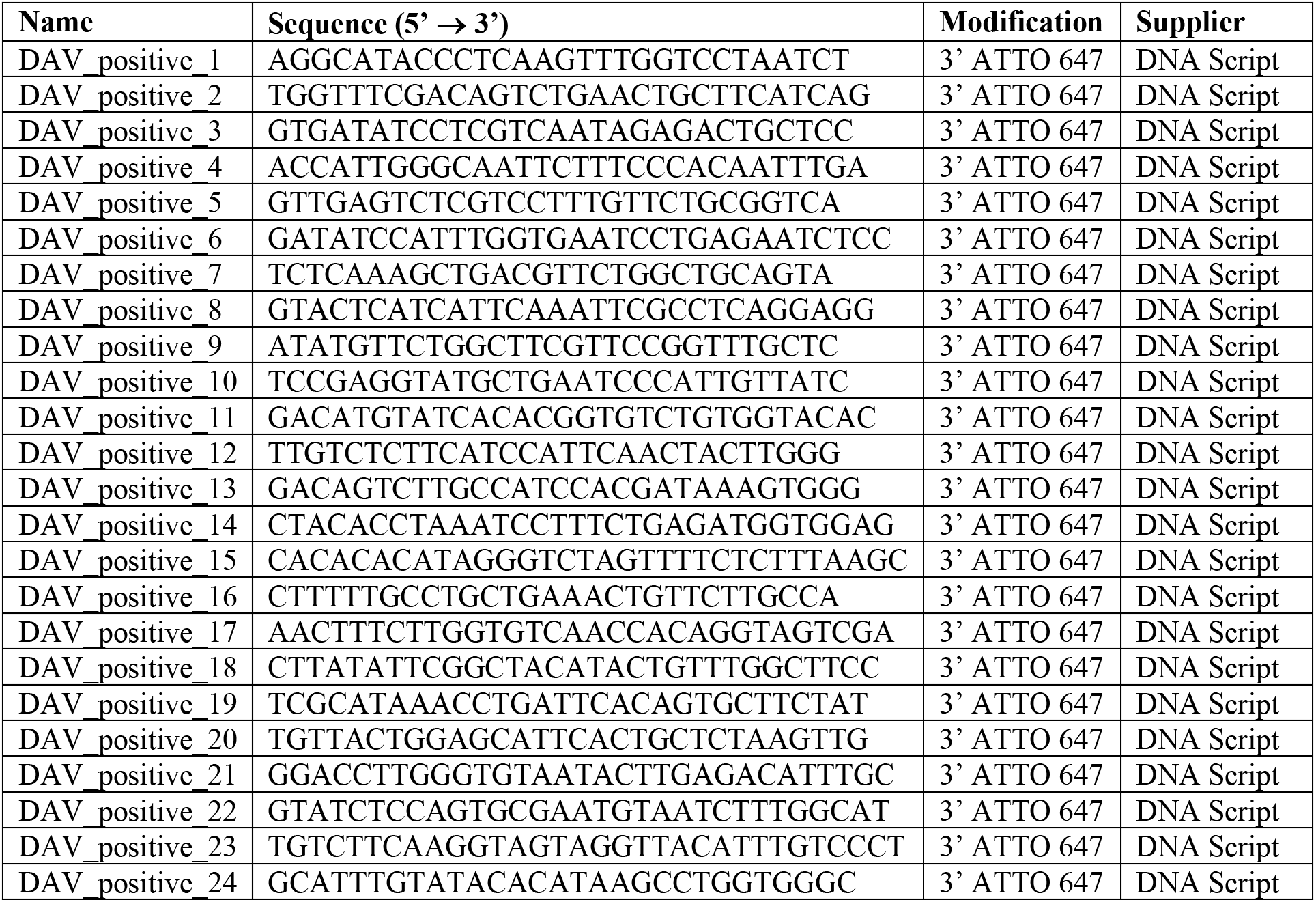
oligonucleotides used to detect positive strand DAV RNA by RNA FISH, related to STAR Methods.

